# A Division of Labor in the Recruitment and Topological Organization of a Bacterial Morphogenic Complex

**DOI:** 10.1101/847335

**Authors:** Paul D. Caccamo, Maxime Jacq, Michael S. VanNieuwenhze, Yves V. Brun

**Affiliations:** Department of Biology, Indiana University, 1001 E. 3rd St, Bloomington, IN 47405, USA; School of Life Sciences, Biodesign Center for Mechanisms of Evolution, Arizona State University, Tempe, Arizona, USA; Département de Microbiologie, Infectiologie et Immunologie, Université de Montréal, Pavillon Roger-Gaudry, C.P. 6128, Succursale Centreville, Montréal, Canada; Department of Molecular and Cellular Biochemistry, 212 S. Hawthorne Drive, Indiana University, Bloomington, IN 47405, USA; Department of Chemistry, Indiana University, 800 East Kirkwood Avenue, Bloomington, IN 47405, USA

## Abstract

Bacteria come in an array of shapes and sizes, but the mechanisms underlying diverse morphologies are poorly understood. The peptidoglycan (PG) cell wall is the primary determinant of cell shape. At the molecular level, morphological variation often results from the regulation of enzymes involved in cell elongation and division. These enzymes are spatially controlled by cytoskeletal scaffolding proteins, that both recruit and organize the PG synthesis complex. How then do cells define alternative morphogenic processes that are distinct from cell elongation and division? To address this, we have turned to the specific morphotype of Alphaproteobacterial stalks. Stalk synthesis is a specialized form of zonal growth, which requires PG synthesis in a spatially constrained zone to extend a thin cylindrical projection of the cell envelope. The morphogen SpmX defines the site of stalk PG synthesis, but SpmX is a PG hydrolase. How then does a non-cytoskeletal protein, SpmX, define and constrain PG synthesis to form stalks? Here we report that SpmX and the bactofilin BacA act in concert to regulate stalk synthesis in *Asticcacaulis biprosthecum*. We show that SpmX recruits BacA to the site of stalk synthesis. BacA then serves as a stalk-specific topological organizer for PG synthesis activity, including its recruiter SpmX, at the base of the stalk. In the absence of BacA, cells produce “pseudostalks” that are the result of unconstrained PG synthesis. Therefore, the protein responsible for recruitment of a morphogenic PG remodeling complex, SpmX, is distinct from the protein that topologically organizes the complex, BacA.

## Introduction

Bacterial cells come in a panoply of shapes and sizes, from the ubiquitous rods and cocci to hyphal, star-shaped, and appendaged bacteria [1]. In addition to shapes that are reproduced faithfully across generations, bacterial cells can dynamically change shape in response to environmental conditions or through a programmed life cycle [1–3]. The shape of most bacteria is determined by the peptidoglycan (PG) cell wall, and the resulting morphology is the product of complex interactions between the proteins and regulatory elements that compose the PG biosynthetic machinery. Much morphological variation results from the differential regulation of divisome or elongasome proteins. For example, the suppression of divisome proteins allows ovococcoid bacteria to form rods under certain biofilm conditions or pathogenic rod-shaped bacteria to filament during infection [3–6]. Filamentous Streptomycetes form branching hyphae by localizing the same elongation machinery in different places [3, 7]. How then do cells define a new morphological process that is distinct from the elongasome or divisome, but still operates in the context of those critical PG synthesis modes? To address this question, we have turned to the specific morphotype of Alphaproteobacterial prosthecae.

Prosthecae, or “stalks”, are non-essential extensions of the bacterial cell body thought to play a role in nutrient uptake [8, 9]. Stalk synthesis requires PG synthesis in a spatially constrained zone in order to extend a thin cylindrical projection of the cell envelope. The stalk is compartmentalized from the cytoplasm, as it is devoid of DNA and ribosomes and excludes even small cytoplasmic proteins such as GFP [9]. *Asticcacaulis biprosthecum*, a gram negative Alphaproteobacterium from the Caulobacteraceae family [10], produces two bilateral stalks whose synthesis depends on PG synthesis at the base of the incipient stalk structures (red arrows in Figure 1A) [11]. How do cells harness PG synthesis machinery to produce stalks while preventing its typical function of cell elongation or division?

**Figure 1.**
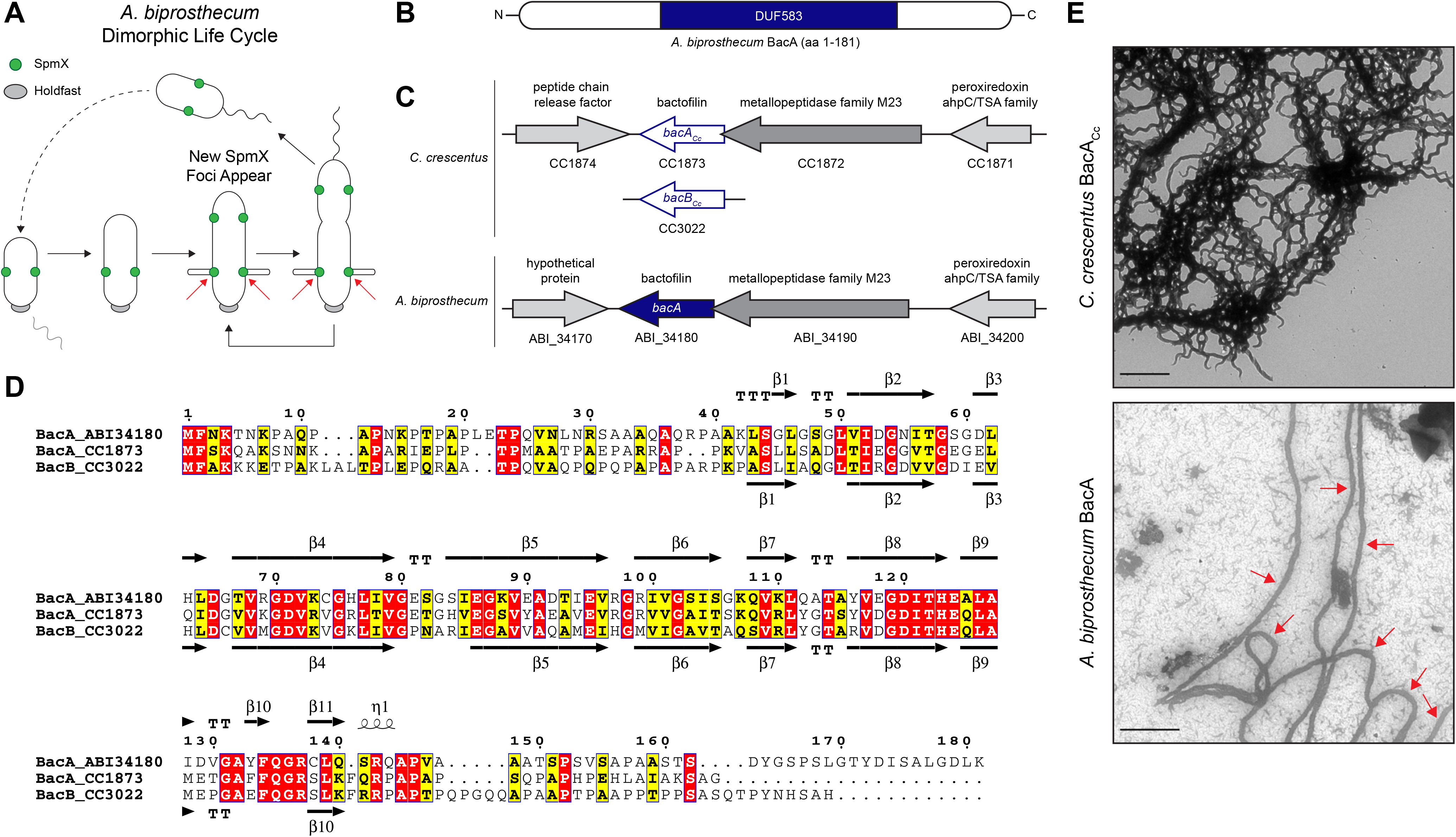
Comparison of *Caulobacter crescentus* and *Asticcacaulis biprosthecum* Bactofilins. (A) Dimorphic life cycle of *A. biprosthecum*. A common characteristic of prosthecate Alphaproteobacteria, the prosthecate mother cell produces an adhesive holdfast (shown in grey) at one pole. Cell division results in a motile, nonreplicating swarmer cell that differentiates into a prosthecate cell. Red arrows indicate the base of *A. biprosthecum* stalks. Green circles indicate SpmX localization. (B) Schematic of the BacA protein from *A. biprosthecum*. The conserved bactofilin (DUF583) domain is shown in blue. (C) Schematics of the bactofilin gene loci for both *C. crescentus* and *A. biprosthecum*. (D) Alignment of *A. biprosthecum* BacA (ABI_34180) and *C. crescentus* BacA_Cc_/BacB_Cc_ (CC1873/CC3022). Conserved residues are shown in red boxes, similar residues in yellow boxes with bold characters. The secondary structure of *A. biprosthecum* BacA (“BacA_ABI34180”) was modeled using the SWISS-MODEL server [50] with the structure of *C. crescentus* BacA_Cc_ (“BacA_1873”; PBD code 2N3D) as a template [51]. The secondary structures of *C. crescentus* BacA_Cc_ (PDB code 2N3D) and the predicted ones for *A. biprosthecum* BacA are indicated below and above the sequence alignment, respectively. Residues are numbered according to *A. biprosthecum* BacA. (E) Transmission electron microscopy (TEM) of BacA protofilament bundles from *C. crescentus* taken in detergent free conditions (top) and *A. biprosthecum* taken in the presence of 10% Triton X-100 (bottom). To distinguish from background, red arrows indicate *A. biprosthecum* BacA protofilaments. Scale bars = 1 μm. See also Table S1.

Here we report how a recently identified class of bacterial cytoskeletal protein, known as “bactofilin” (BacA), plays a dual role by defining the topography of PG synthesis for the synthesis of stalks and by inhibiting a default cell elongation mode at that same site in *A. biprosthecum*. Bactofilins are conserved throughout the bacterial kingdom and are characterized by the presence of a central conserved DUF583 (or “bactofilin”) domain flanked by N- and C-terminal regions of variable length and sequence (Figure 1B) [12]. Bactofilins are involved in cell shape determination in a number of species. For example, in the helical *Helicobacter pylori* and *Leptospira biflexa*, bactofilins are required for proper helical shape generation [13, 14]. In *Caulobacter crescentus*, bactofilins optimize the rate of stalk synthesis at the cell pole by a yet unknown mechanism [12]. A bactofilin mutant of *C. crescentus* still synthesizes stalks normally in a complex medium and exhibits a 50% reduction in stalk length under phosphate starvation where stalk elongation is strongly stimulated. While bactofilin recruits PbpC, whose mutant leads to a similar modest reduction in stalk length, a bactofilin mutant still exhibits strong PG synthesis activity similar to wild-type cells, indicating that *C. crescentus* bactofilin is not required for stalk PG synthesis. Therefore, it is not well understood how bactofilins exert an effect on morphology, but a common theme appears to be association or direct interaction with PG modifying enzymes.

The best studied bacterial cytoskeletal proteins (MreB, FtsZ, and DivIVA) serve dual roles as both recruiters and organizers of the proteins for their respective PG synthesis complexes. In contrast, we show that the roles of recruiter and organizer of PG synthesis have undergone a division of labor in *A. biprosthecum* stalk synthesis, where a morphogen, SpmX, recruits its own organizing cytoskeletal protein, BacA. Our prior work established that the recruitment of the stalk PG synthesis machinery is performed by the modular morphogen SpmX: changing SpmX location is sufficient to drive stalk synthesis at a new site [11], much like changing the position of FtsZ changes the location of the site of cell division. However, SpmX is a PG hydrolase, not a cytoskeletal protein [11, 15]. How then does a non-cytoskeletal protein, SpmX, define and constrain PG synthesis to form stalks? In this work, we identify BacA as a cytoskeletal scaffolding protein that provides topological specificity to stalk PG synthesis in *A. biprosthecum*, including to the recruiter SpmX itself. We apply genetics, cell biology, microscopy, chemical labeling, and quantitative analysis to examine the interconnected roles of the morphogen SpmX and the bactofilin BacA in *A. biprosthecum* stalk synthesis. We present evidence that SpmX and BacA act in a coordinated fashion to initiate and regulate stalk synthesis. SpmX acts as the recruiter of the stalk synthesis complex, including BacA. BacA then serves as a stalk-specific cytoskeletal scaffolding protein that anchors the putative synthesis complex, including SpmX, to the base of the stalk, and defines the width of the complex. In the absence of BacA, cells produce abnormal “pseudostalks” that are the result of unconstrained PG synthesis correlated with the mislocalization of SpmX. Finally, we show that BacA is required to block PG synthesis involved in cell elongation and division, as well as DNA, from the site of stalk synthesis defined by SpmX. Therefore, the protein responsible for recruitment of a morphogenic PG remodeling complex, SpmX, is different than the protein that topologically organizes the complex, BacA. Such separation of the recruitment and scaffolding activities in two different proteins may prove more versatile for morphogenetic events that do not simply involve modulation of cell elongation or division.

## Results

### *A. biprosthecum* encodes a single bactofilin gene whose product self-polymerizes *in vitro*

*A. biprosthecum* has proven to be a better system than *C. crescentus* to identify genes required for stalk synthesis because: 1) with the exception of a *C. crescentus* MreB-mCherry sandwich fusion that acts through an unknown mechanism to block stalk synthesis [16], the *A. biprosthecum spmX* mutant is the only known stalkless mutant in any Caulobacterales species whose phenotype is not bypassed by phosphate starvation [11, 17, 18], which stimulates stalk synthesis [11, 19–21]; 2) *A. biprosthecum* stalks are located at midcell, away from the myriad of polarly localized proteins involved in *C. crescentus* development and cell cycle progression [22–25], eliminating any potentially confounding co-localization or interaction data; and 3) many of the genetic tools developed for *C. crescentus* work in *A. biprosthecum*. In *C. crescentus*, deletion of the bactofilin genes *bacA* and *bacB* (for clarity, these will be referred to as “*bacA*_*Cc*_” and “*bacB*_*Cc*_”) leads to a slight reduction in the length of otherwise normal stalks [12, 26, 27]. Only one bactofilin homolog was found in the *A. biprosthecum* genome, which we have named “*bacA*” due to its bidirectional best hit in *C. crescentus* being *bacA*_*Cc*_ (Figure 1C). The genomic arrangement of the *A. biprosthecum bacA* locus is similar to that of *C*. *crescentus bacA*_*Cc*_, with a putative M23 family metallopeptidase gene (CC1872/ABI_34190) directly upstream and overlapping the *bacA_Cc_/bacA* coding regions, and a putative peroxiredoxin gene (CC1871/ABI_34200) further upstream (Figure 1C). Downstream of and convergent to *bacA* lies a hypothetical gene (Figure 1C). BacA exhibits high sequence and predicted structural similarity to the *C. crescentus* bactofilins, with a central DUF583 domain composed of repeating β strands flanked by proline-rich N- and C-termini, suggesting similar biochemical properties (Figure 1D).

Indeed, BacA_Cc_ can self-assemble *in vitro* into filaments, filament bundles, and sheets [12, 28]. To test if *A. biprosthecum* BacA is a self-polymerizing protein, we overexpressed and purified both BacA_Cc_ and BacA in *Escherichia coli* and visualized the purified proteins via transmission electron microscopy (TEM) (Figure 1E). While BacA_Cc_ readily polymerized in various environments (salt, pH, chaotropic agent), under similar conditions BacA formed proteins aggregates instead of filaments. While most conditions tested did not improve BacA solubility, addition of high detergent concentration (10% Triton X-100) allowed filament formation. The TEM micrographs show that both bactofilin proteins, BacA_Cc_ and BacA, self-polymerize *in vitro* to form protofilament bundles, with BacA forming ~90 nm wide filaments (Figure 1E; red arrows indicate BacA protofilaments).

### BacA is required for stalk synthesis

In *C. crescentus*, deletion of the bactofilin homologs *bacA*_*Cc*_ and/or *bacB*_*Cc*_ does not abrogate stalk synthesis or impact stalk ultrastructure, but results in a slight reduction in stalk length under stalk synthesis-stimulating, phosphate-limited conditions, and does not impact the strong PG synthesis activity at the base of the stalk [12]. This suggests that, in *C. crescentus*, deletion of *bacAB*_*Cc*_ affects only stalk longitudinal extension and to a limited extent [12]. Strikingly, ~95% of *A. biprosthecum* Δ*bacA* mutant cells were completely stalkless when grown in the rich medium PYE (see culturing details in **Experimental Model and Subject Details**), with a small proportion of cells exhibiting bump-like protrusions at the midcell where stalks would normally be found (Figure S1A). Since growth in rich medium may mask latent stalk phenotypes [11, 21], the Δ*bacA* mutant was studied under phosphate starvation, which stimulates stalk synthesis [11, 19–21].

In low phosphate medium, more than half (58% ± 12%) of WT cells possessed a visible cell body extension, all of which were stalks (Figure 2). In the Δ*bacA* mutant, only 3% ± 2% of total cells produced a stalk (Figure 2). Instead of synthesizing wildtype stalks, the Δ*bacA* mutant produced short, wide protrusions at the bilateral positions where stalks are normally synthesized, which we have termed “pseudostalks” (Figure 2A and red arrows in Figure 2B). Under phosphate starvation, the stalks produced by WT cells were 7.3 ± 3.7 μm long and 171 ± 13 nm in diameter at the base (Figure 2). Pseudostalks were significantly shorter than stalks at 0.9 ± 0.5 μm long (~88% decrease in length compared to WT), were more variable in diameter, and were often branched or “frazzled” at the ends (Figure 2). Pseudostalks were significantly wider in diameter at the base, 392 ± 73 nm or an ~129% increase in width compared to WT (Figure 2).

**Figure 2.**
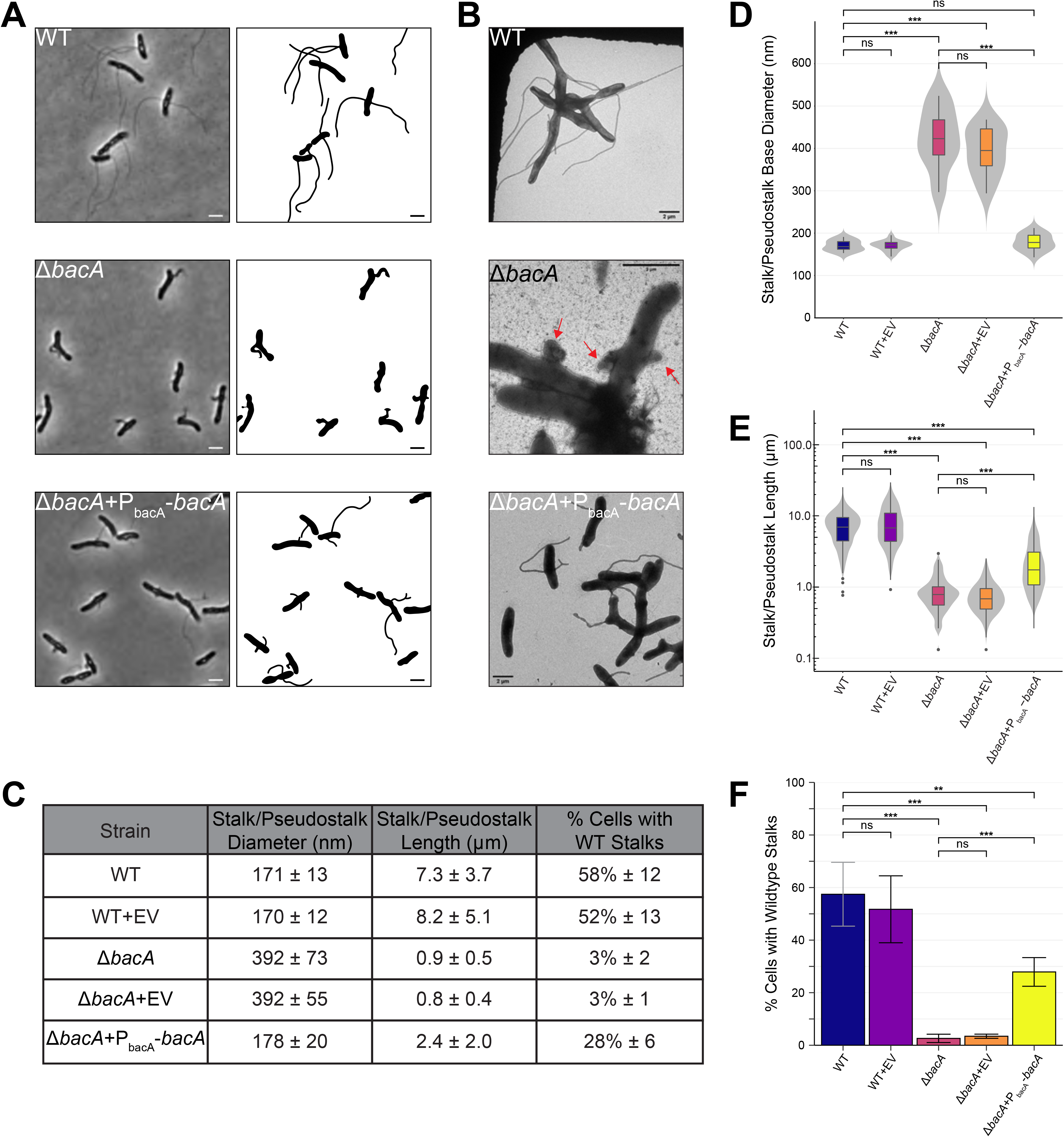
BacA is Required for WT Stalk Synthesis. Analysis of stalk morphology in strains YB642 (WT), YB9183 (WT + pMR10 empty vector control), YB8597 (Δ*bacA*), YB8620 (Δ*bacA* + pMR10 empty vector control), and YB8601 (Δ*bacA* + pMR10-P_bacA_-*bacA*). Cells were grown in rich medium (PYE) to saturation and sub-cultured into phosphate limited (HIGG) medium at 26°C for 72h (see culturing details in Experimental Model and Subject Details). (A) Phase microscopy (left) and cell silhouette (right) for strains YB642 (WT), YB8597 (Δ*bacA*), and YB8601 (Δ*bacA* + pMR10-P_bacA_-*bacA*). Scale bars = 2 μm. (B) Transmission electron microscopy (TEM) for strains YB642 (WT), YB8597 (Δ*bacA*), and YB8601 (Δ*bacA*+pMR10-P_bacA_-*bacA*). Δ*bacA* pseudostalks are indicated with red arrows. Scale bars = 2 μm. (C) Summary statistics for data presented in Figure 2D, 2E, and 2F. Data shown is mean (SD). (D) Distribution of stalk/pseudostalk base diameter in the populations. Data (WT n=13; WT+EV n=19; Δ*bacA* n=8; Δ*bacA*+EV n=13; Δ*bacA*+P_bacA_-*bacA* n= 17) are from single samples with five TEM fields per sample. Data are represented as box and whisker plots [52] where the middle line represents the median, the lower and upper hinges correspond to the first and third quartiles (the 25th and 75th percentiles), and the upper whisker extends from the hinge to the largest value no further than 1.5 * IQR from the hinge (where IQR is the inter-quartile range, or distance between the first and third quartiles). The lower whisker extends from the hinge to the smallest value at most 1.5 * IQR of the hinge. Data beyond the end of the whiskers are called “outlying” points and are plotted individually. A mirrored density violin plot [53] is underlaid to show the continuous distribution of the data. Violin plots have been scaled to the same width. (***p ≤ 0.001, ns = not significant; two-sided t-test). (E) Distribution of stalk/pseudostalk lengths in the populations, measured from the tip of the structure to the cell body. Data (WT n=412; WT+EV n=321; Δ*bacA* n=153; Δ*bacA*+EV n=148; Δ*bacA*+P_bacA_-*bacA* n=324) are from three independent biological replicates with five phase microscopy fields per replicate. Data are plotted on a log10 scale y-axis and are represented as box and whisker plots and violin plots in the same manner as Figure 2D. As length data are approximately log-normally distributed, significant difference testing was performed on log10 transformed data (***p ≤ 0.001, ns = not significant; two-sided t-test). (F) Percentage of cells with WT stalks. Cells in phase images from Figure 2A were scored as having a WT stalk (i.e. a thin extension from the cell body). Cells exhibiting thick and aberrant pseudostalks were excluded. Data (total cells counted: WT n=735; WT+EV n=615; Δ*bacA* n=841; Δ*bacA*+EV n=1259; Δ*bacA*+P_bacA_-*bacA* n=831) are from four independent biological replicates with five phase microscopy fields per replicate. Data are represented as the mean (SD) percentage of cells with stalks. (***p ≤ 0.001, **p ≤ 0.01, ns = not significant > 0.05; two-sided t-test). See also Figure S1.

The presence of a low-copy plasmid encoding *bacA* expressed from its native promoter restored stalk diameter to WT levels (178 ± 20 nm vs. 171 ± 13 nm, respectively), suggesting that BacA plays a key role in determining and maintaining stalk width (Figure 2). In addition, 28% ± 6% of complemented Δ*bacA* cells produced WT-like stalks, a 10-fold increase over the mutant (Figure 2). Complemented Δ*bacA* cells produced stalks 2.4 ± 2.0 μm long that, while shorter than WT stalks, were significantly longer than Δ*bacA* pseudostalks (Figure 2). It is unlikely that the less efficient complementation of stalk number and length are due to polarity effects of the mutation since the gene downstream of *bacA* is transcribed in the opposite direction (Figure 1C). Dosage effects may explain why complementation is not fully achieved for these phenotypes; however, we are unable to test this at this time due to a lack of BacA antibodies. Taken together, these data show that BacA is required for stalk synthesis in *A. biprosthecum*.

### BacA localizes to the site of stalk synthesis after SpmX localization

While the localization of SpmX is known, the pattern of BacA localization is not. The stalk morphogen SpmX localizes to the future site of stalk synthesis in predivisional cells of *Asticcacaulis*, where it marks the site of stalk synthesis, which occurs after cell division and swarmer cell differentiation in the next cell cycle (Figure 1A) [11]. To investigate the subcellular localization of BacA in WT cells, we fused BacA to mVenus at the native locus and performed fluorescence microscopy. This fusion is functional, and cells expressing BacA-mVenus produced normal stalks at a similar frequency as WT cells expressing the native protein (Figure S1).

We first performed time-lapse fluorescence microscopy to determine the timing of BacA localization during the cell cycle (Figure 3A and Videos S1 & S2). The first four panels of Figure 3A (0-84 min) show the elongation of a predivisional mother cell as it produces an incipient swarmer daughter cell, where BacA is already localized at bilateral positions in the stalked half of the cell. In contrast to SpmX, which localizes before cell division at the future sites of stalk synthesis, we were unable to detect new BacA foci in predivisional cells. After cell division (the transition from 84-112 min), a BacA-mVenus focus appeared at a lateral position in the incipient swarmer cell. While only one BacA-mVenus focus is visible in the overlay from 112 to 140 min, a second focus is clearly visible in the overlay at 168 and 196 min. This process of BacA localization after cell division is repeated in later panels with a second daughter cell (196-252 min). Therefore, SpmX localizes before cell division (Figure 4A), whereas BacA localizes after cell division, significantly after SpmX (Figure 3C). We then sought to quantify BacA-mVenus localization at the population level, rather than single cells. Overall, BacA-mVenus localized to bilateral positions at the base of the stalks (Figure 3D). A population-level heatmap of the subcellular localization of BacA-mVenus foci exhibited a localization pattern similar to that of SpmX (Figure 4D) [11], with foci clustering in a bilateral manner at the midcell (Figure 3E).

**Figure 3.**
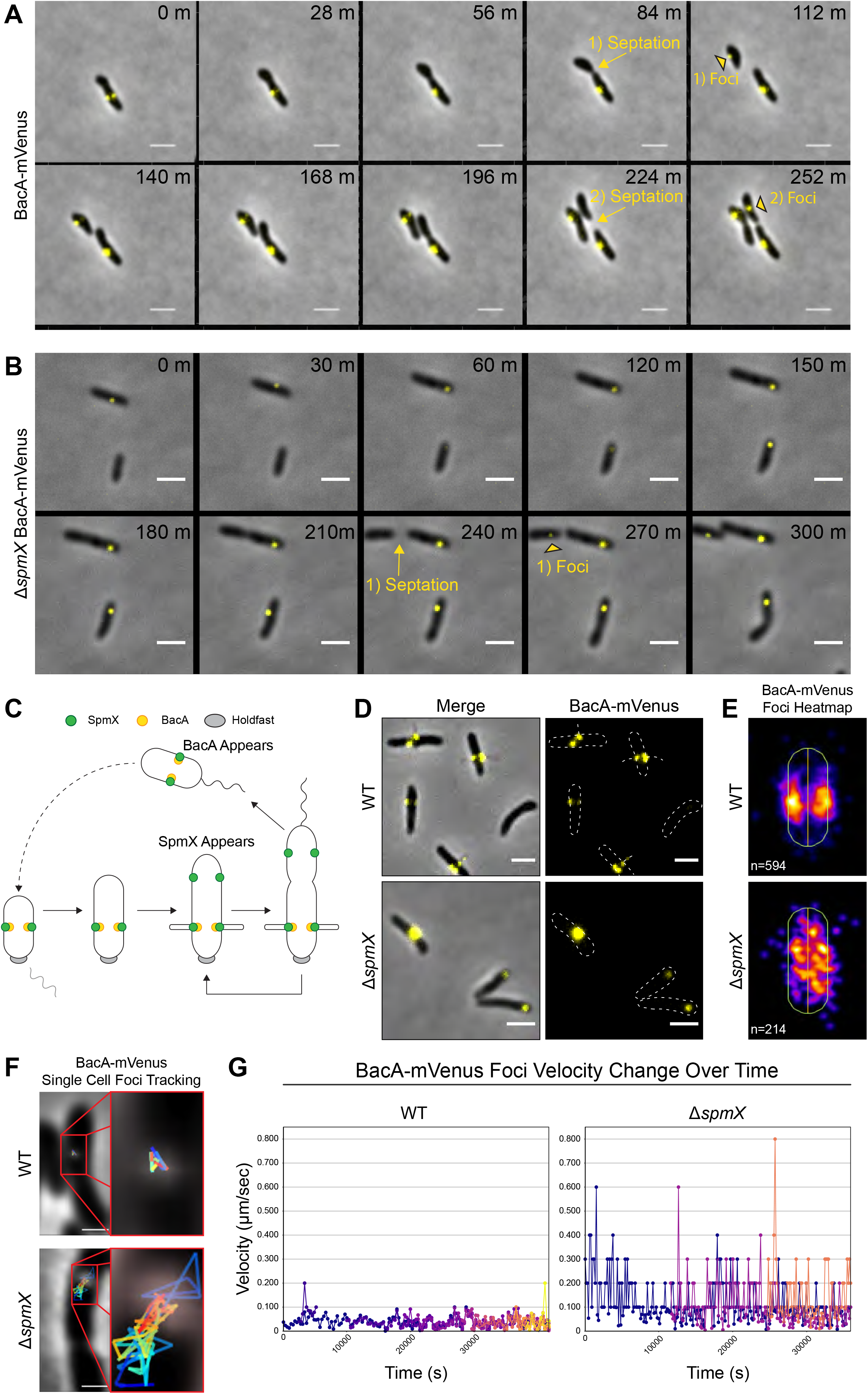
BacA Localizes After Cell Division, Localizes to the Base of Stalks, and Requires SpmX for Localization. Subcellular localization of BacA-mVenus in strains YB7474 (*bacA*::*bacA-mVenus*) and YB7487 (Δ*spmX bacA*::*bacA-mVenus*) grown in rich medium (PYE) at 26°C for 24-48h. (A-B) Time lapse microscopy montage showing the dynamics of BacA-mVenus localization in (A) strain YB7474 (*bacA*::*bacA-mVenus*) or (B) strain YB7487 (*bacA*::*bacA-mVenus* Δ*spmX*). Yellow arrows and text show the transition from pre-septation to post-septation. Yellow black-bordered triangles mark the appearance of new BacA-mVenus foci in recently divided daughter cells. Septation events and the corresponding foci appearance are numbered. Frames show images taken every (A) 28 minutes or (B) 30 minutes. Scale bars = 2 μm. (C) Dimorphic life cycle of *A. biprosthecum*. The prosthecate mother cell produces an adhesive holdfast (shown in gray) at one pole. Cell division results in a motile, nonreplicating swarmer cell that differentiates into a prosthecate cell. Green circles indicate SpmX localization. Yellow circles indicate BacA localization. (D) BacA-mVenus subcellular localization. Phase merge and fluorescence microscopy images. Scale bars = 2 μm. (E) Population level heatmaps of BacA-mVenus subcellular localization. Number of foci analyzed is shown on the bottom left of each heatmap. (F) Single cell BacA-mVenus foci tracking. Tracks are colored by time, with blue representing the earliest timepoints and red representing the latest time points. Scale bars = 1 μm. (G) Velocity tracking of multiple BacA-mVenus foci from time-lapse videos. Particle tracking was performed and velocity (μm/s) from frame-to-frame for each focus was calculated. Each color represents the track of individual foci as they appear. See also Figures S1 & S3 and Videos S1-S4.

**Figure 4.**
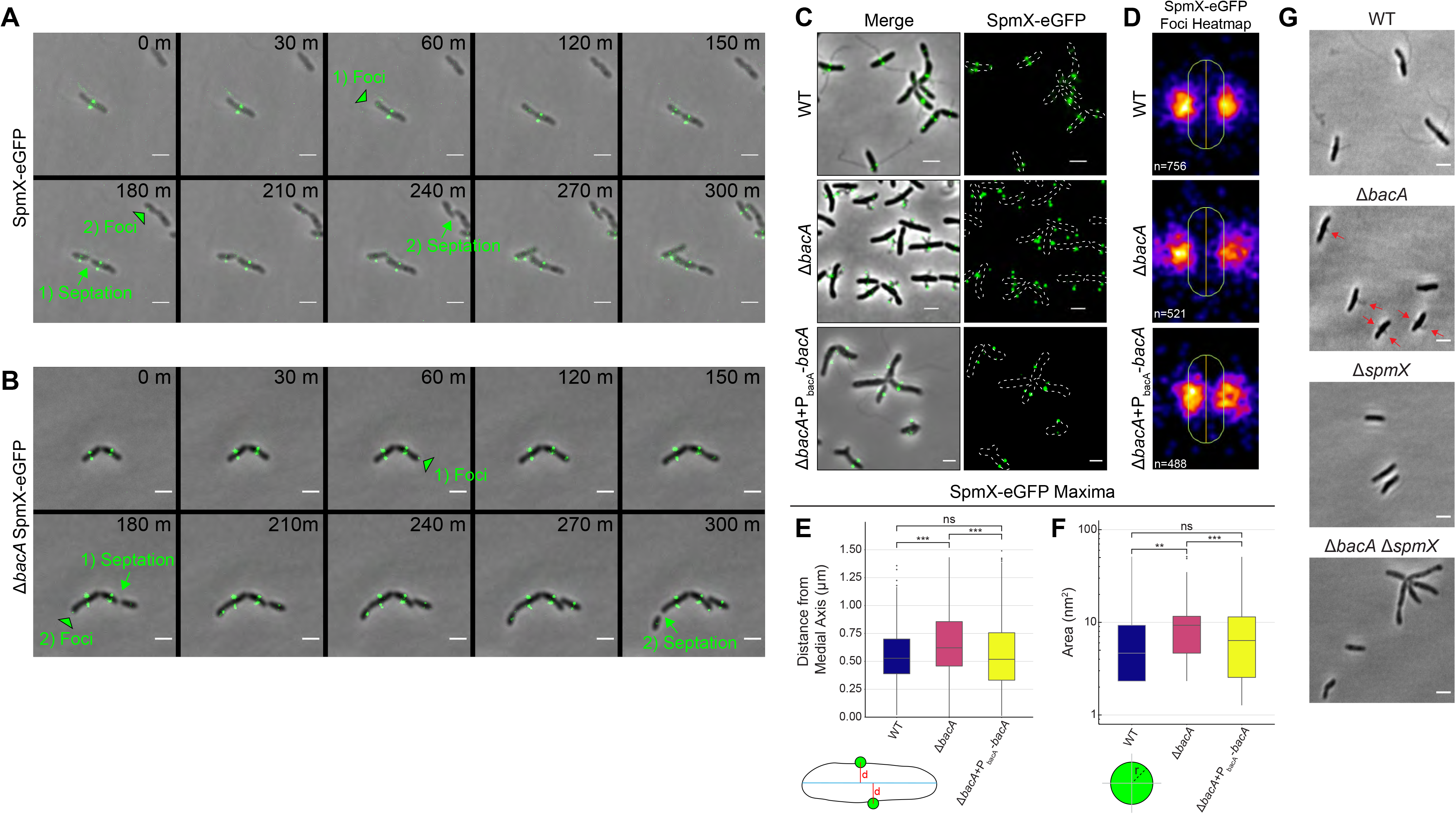
BacA is not Required to Localize SpmX-eGFP, but is Required to Constrain SpmX-eGFP at the Base of the Stalk and to Constrain the Size of SpmX-eGFP Maxima. (A-F) Subcellular localization of SpmX-eGFP in strains YB5692 (*spmX*::*spmX-eGFP*), YB7561 (*spmX*::*spmX-eGFP* Δ*bacA*), and YB9521 (*spmX*::*spmX-eGFP* Δ*bacA+*P_bacA_-*bacA*). Cells were grown in rich medium (PYE) to saturation and sub-cultured into phosphate limited (HIGG) medium at 26°C for 72h. (A-B) Time lapse microscopy montage showing the dynamics of SpmX-eGFP localization in (A) strain YB5692 (*spmX*::*spmX-eGFP*) or (B) strain YB7561 (*spmX*::*spmX-eGFP* Δ*bacA*). Green black-bordered triangles mark the appearance of new SpmX-eGFP foci in predivisional daughter cells. Green arrows and text show the transition from pre-septation to post-septation. Foci appearance and the corresponding septation events are numbered. Frames show images taken every 30 minutes. Scale bars = 2 μm. (C) SpmX-eGFP subcellular localization. Phase merge and fluorescence microscopy images. Scale bars = 2 μm. (D) Population level heatmaps of SpmX-eGFP subcellular localization. Number of foci analyzed is shown on the bottom left of each heatmap. (E) Distribution of orthogonal distance (μm) of each SpmX-eGFP maxima from the medial axis of its associated parent cell. Data set is the same as used in Figure 4D. Data are plotted on a continuous y-axis and are represented as box and whisker plots as described in Figure 2D. (***p ≤ 0.001, ns = not significant > 0.05; two-sided t-test). (F) Distribution of SpmX-eGFP maxima area (nm^2^) in the populations. Data set is the same as used in Figure 4D. Data are plotted on a log_10_ scale y-axis and are represented as box and whisker plots as described in Figure 2D. (***p ≤ 0.001, **p ≤ 0.01, ns = not significant > 0.05; two-sided t-test). (G) A Δ*bacA* Δ*spmX* double mutant phenocopies a Δ*spmX* single mutant. Representative phase images of YB642 (WT), YB8597 (Δ*bacA*), YB8237 (Δ*spmX*), and YB7489 (Δ*bacA* Δ*spmX*). Cells were grown in rich medium (PYE) to saturation and sub-cultured into phosphate limited (HIGG) medium at 26°C for 48h-72h before imaging. WT cells produce stalks (top left), Δ*bacA* cells produce pseudostalks (top right, red arrows), and both Δ*spmX* and Δ*bacA* Δ*spmX* cells are stalkless (bottom). Scale bars = 2 μm. See also Figures S2 & S3 and Videos S5 & S6.

### SpmX is required for BacA localization

Considering that 1) both SpmX and BacA are required for WT stalk synthesis; 2) that both proteins localize to the site of stalk synthesis; and 3) that SpmX localizes to the site of stalk synthesis prior to BacA, we hypothesized that SpmX is required for BacA localization. In order to test the role of SpmX in BacA localization, we constructed a strain with a *bacA*-*mVenus* fusion at the native locus in the Δ*spmX* background.

As reported above in the WT background, BacA-mVenus localized to the base of the stalk (Figure 3D, top) with foci clustering in a bilateral manner at the midcell (Figure 3E, top). In the Δ*spmX* mutant, BacA-mVenus was often mislocalized toward the poles (Figure 3D, bottom), and the population-level heatmap of the subcellular localization of BacA-mVenus foci in the Δ*spmX* mutant showed that BacA-mVenus was randomly distributed throughout the cell body (Figure 3E, bottom). However, these images represent only a snapshot of BacA-mVenus localization at a single time point. To further investigate BacA-mVenus dynamics in WT and Δ*spmX* strains, we performed time-lapse fluorescence microscopy (Figures 3A & 3B and Videos S1-S4) and used particle tracking to follow movement and velocity changes for individual foci (Figures 3F & 3G). In WT cells, BacA-mVenus foci were constrained bilaterally at sites of stalk synthesis and remained relatively static once they appeared (Figures 3A, 3F, & 3G and Videos S1 & S2). Conversely, in the Δ*spmX* mutant, BacA-mVenus foci moved randomly throughout the cell body and exhibited start-stop patterns of spikes and drops in velocity (Figures 3B, 3F, & 3G and Videos S3 & S4). Taken together, these data show that SpmX is required for BacA localization.

### BacA is required to constrain SpmX at the base of the stalk

Cytoskeletal scaffolding proteins can be loosely defined as proteins that physically organize the molecular components of a biological process or pathway [3]. Because 1) SpmX acts in a modular fashion to define the site of stalk synthesis [11]; 2) SpmX is required for BacA localization; and 3) BacA is required to constrain stalk diameter, we hypothesized that BacA organizes PG synthesis enzymes, and perhaps SpmX, at the site of stalk synthesis. To test SpmX localization in the absence of BacA, we constructed strains with a *spmX*-*eGFP* fusion at the native locus in both WT and Δ*bacA* backgrounds.

Consistent with previously published results [11], fluorescence microscopy showed that SpmX-eGFP first appears in the incipient swarmer compartment of predivisional cells of *A. biprosthecum* where it localizes bilaterally to mark the site of stalk synthesis, which occurs after cell division and swarmer cell differentiation [11] (Figures 1A, 4A, 4C, & 4D and Video S5). Interestingly, SpmX localized to the future bilateral site of stalk synthesis even in the absence of BacA (Figures 4B-4D and Video S6). However, in the Δ*bacA* mutant SpmX-eGFP often localized to the tips of pseudostalks rather than the base of stalks as it does in WT cells (Figure 4C). When stalk synthesis was restored in the Δ*bacA* mutant by the expression of BacA from a plasmid, SpmX localization to the base of the stalk was restored as well (Figure 4C). This defect of SpmX localization is further revealed by the localization heatmaps, where, rather than a tight localization close to the cell envelope in WT, there is a diffuse localization at the midcell that radiates away from the cell body in the Δ*bacA* mutant (Figure 4D). Indeed, quantification of the orthogonal distance of SpmX-eGFP maxima from the medial axis of the associated cell, showed a significant increase in Δ*bacA* cells compared to WT, a defect that was rescued in the complementation strain (Figure 4E). It should be noted that when stalk synthesis is restored by exogenous expression of BacA, SpmX localization to the base of the stalk is also restored in these cells (Figure S2). While global SpmX protein levels according to Western blot are similar between WT and Δ*bacA* strains (Figure S3), the area of the SpmX-eGFP maxima significantly increased in the Δ*bacA* mutant compared to WT, a defect that was restored in the complementation strain (Figure 4F). This suggests that BacA may play a role in regulating the number of SpmX molecules that comprise the stalk synthesis complex. These results indicate a scaffolding role for BacA during stalk synthesis whereby BacA is required to constrain SpmX in a tight area at the base of the stalk. The above results suggest that SpmX acts upstream of BacA, which was confirmed by the Δ*spmX* Δ*bacA* double mutant, which phenocopied *ΔspmX* (Figure 4G).

### BacA and SpmX colocalize *in vivo* and interact in a bacterial two-hybrid system

Since SpmX is required for BacA localization (Figure 3) and BacA is required to constrain SpmX at the base of the stalk (Figure 4), we wondered whether this could be due to a direct interaction between the two proteins. To address this, we first simultaneously expressed SpmX-mCherry and BacA-mVenus fusions in *A. biprosthecum*. Cells produced stalks, indicating that the fusions are functional when expressed together. Both BacA-mVenus and SpmX-mCherry formed foci at the midcell that often overlapped (Figure 5A), and there was a positive correlation between the localization of the two fusion proteins (Pearson’s Correlation Coefficient = 0.48 ± 0.23; n = 179). We note that in many cases, one of the co-localized BacA focus was much weaker than the other, with some cells having only detectable co-localized BacA with one of the two SpmX foci. This may be an indication that relatively few molecules of BacA are required to constrain SpmX at the base of stalks or that the BacA-SpmX interaction is transitory and becomes less important as the stalk synthesis matures.

**Figure 5.**
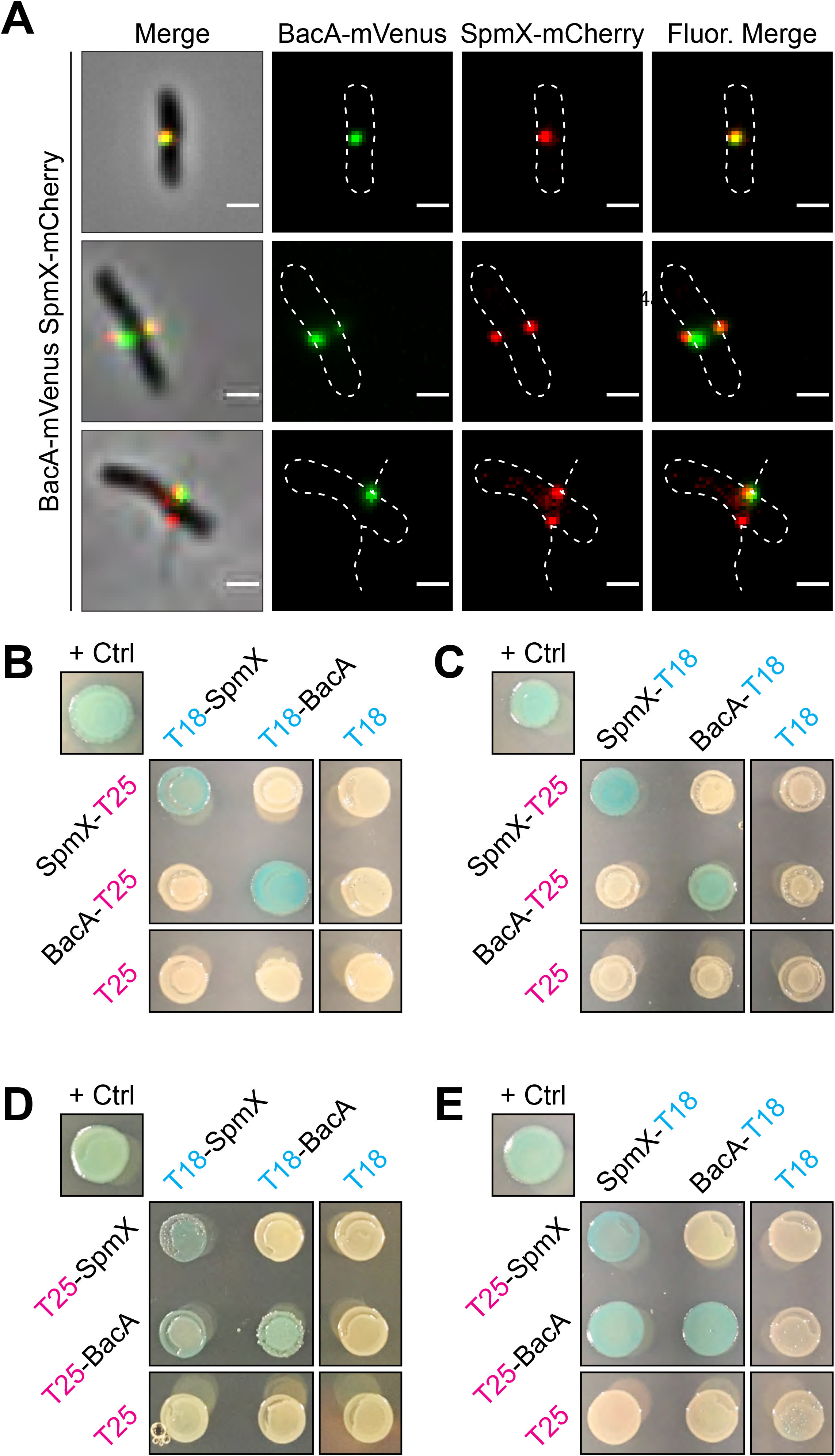
BacA-mVenus and SpmX-mCherry *in vivo* Colocalization and Bacteria Adenylate Cyclase Two-Hybrid (BACTH) Assays Showing *in vivo* Interaction of SpmX and BacA. (A) Representative images of dual labeled strain YB9466 (*bacA*::*bacA-mVenus spmX*::*spmX-mCherry*), with a cell exhibiting almost complete overlap of signals (top), a cell exhibiting BacA-mVenus and SpmX-mCherry foci at both putative sites of stalk synthesis (middle), and a cell with WT stalks, showing functionality of the fusion proteins (bottom). Pearson’s Correlation Coefficient (PCC) was used to quantify colocalization of signal intensity between the two fluorophore channels (BacA-mVenus and SpmX-mCherry) within the cells. PCC values range from −1 indicating a strong negative correlation (anticolocalization) to +1 indicating a strong positive correlation (colocalization), with a value of 0 indicating no correlation (noncolocalization). Only cells that contained both BacA-mVenus and SpmX-mCherry foci located at the midcell were used (n = 179). Mean (SD) PCC = 0.48 ± 0.23, showing an overall positive correlation. (B-E) BTH101 *E. coli* were co-transformed with T25 and T18 based recombinant plasmids. Transformants were then grown in selective medium containing 0.5 mM IPTG and patched on selective indicator plates containing X-gal 40 μg ml^−1^ and 0.5 mM IPTG for ~24 hours at 30°C. As this is a β-galactosidase assay, blue patches indicate a positive *in vivo* interaction between the two recombinant proteins. Positive control (“+ Crtl”) for all assays is T25-Zip and T18-Zip fusion proteins. Negative controls are unfused T18 or T25 fragments in combination with the experimental fusion protein or the cognate unfused T25 or T18 fragments. Patches shown are representative of independent biological triplicates for each combination. (B) C-terminal fused T25 + N-terminal fused T18. (C) C-terminal fused T25 + C-terminal fused T18. (D) N-terminal fused T25 + N-terminal fused T18. (E) N-terminal fused T25 + C-terminal fused T18.

We then asked whether SpmX and BacA interact *in vivo* using <u>b</u>acterial <u>a</u>denylate <u>c</u>yclase <u>t</u>wo-<u>h</u>ybrid (BACTH) assays [29]. We tested all combinations of T18 and T25 fusions for SpmX and BacA (Figure 5B-5E). Serving as an internal control, both SpmX and BacA showed self-interactions (Figure 5B-5E), consistent with previous results showing that *C. crescentus* SpmX forms oligomers [30] and confirming our results showing BacA is a self-polymerizing protein (Figure 1E). Notably, we also observed SpmX-BacA interactions (Figures 5D & 5E), although the only interactions we observed were when BacA was fused with the T25 fragment on the N-terminus (T25-BacA). These results make sense in the context of recent work showing that the N-terminal tail of *Thermus thermophilus* bactofilin is required for membrane binding [31]. If the BacA N-terminal tail is also involved in membrane binding, the fact that we only detect SpmX-BacA interactions when the C-terminal tail is unencumbered by a fusion suggests that the C-terminal region of BacA is involved in protein-protein interactions with SpmX.

### Pseudostalks are composed of PG and formed through dispersed PG synthesis

*A. biprosthecum* stalks extend through the addition of newly synthesized PG at the base of the stalk and are composed of PG throughout [11]. Indeed, isolated PG sacculi maintain the shape of the bacterial cell, including PG-containing stalks [32]. To test if pseudostalks are composed of PG, we first isolated sacculi from *A. biprosthecum* WT and Δ*bacA* cells. As expected, *A. biprosthecum* WT sacculi exhibited long and thin bilateral extensions of stalk PG (Figure 6A). Sacculi purified from Δ*bacA* cells showed sac-like bilateral extensions of PG that mimicked the morphology of pseudostalks (Figure 6A), confirming that pseudostalks are composed of PG.

**Figure 6.**
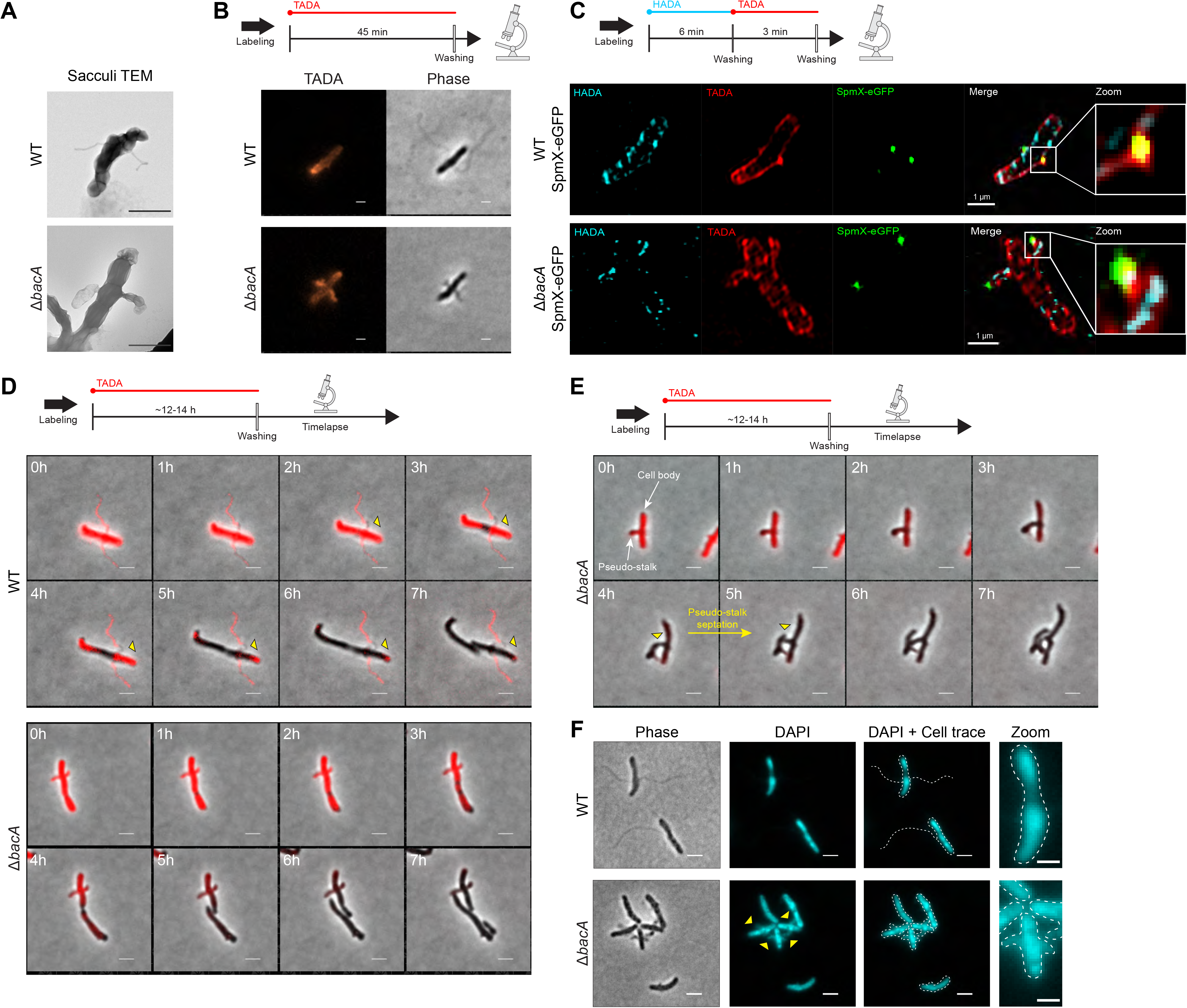
Peptidoglycan (PG) Composition & Remodeling and Septation & DNA Content in WT Stalks and Δ*bacA* Pseudostalks. (A) Transmission electron microscopy (TEM) of prepared PG sacculi for strains YB642 (WT) and YB8597 (Δ*bacA*). Cells were grown in rich medium (PYE) to saturation and sub-cultured into phosphate limited (HIGG) medium at 26°C for 72h before preparing sacculi. Sacculi were prepared by boiling in SDS for 30 minutes followed by washing ≥6X with dH_2_O. Scale bars = 1 μm. (B) Medium pulse FDAA (TADA) labeling showing active PG remodeling in strains YB642 (WT) and YB8597 (Δ*bacA*). Cells were grown in rich medium (PYE) to saturation and sub-cultured into phosphate limited (HIGG) medium at 26°C for 72h) before labeling. Cells were washed with 2X PYE, labeled with 500 μM TADA for 45 minutes, washed 2X with PYE, and imaged with phase and fluorescence microscopy. Representative images are shown. Scale bars = 2 μm. (C) Virtual time lapse: Short pulse, sequential, dual FDAA (TADA and HADA) labeling showing active PG remodeling in strains YB5692 (*spmX*::*spmX-eGFP*) and YB7561 (*spmX*::*spmX-eGFP* Δ*bacA*). Cells were grown in rich medium (PYE) to saturation and sub-cultured into phosphate limited (HIGG) medium at 26°C for 72h before labeling. Cells were washed 2X with PYE, labeled with 500 μM HADA for 6 minutes, washed 2X with PYE, labeled with 500 μM TADA for 3 minutes, washed 2X with PYE, and imaged via 3D-SIM (Structured Illumination Microscopy). Representative images are shown. Scale bars = 1 μm. (D) Pulse-chase time-lapse of FDAA (TADA) labeling for strains YB642 (WT) and YB8597 (Δ*bacA*) showing loss of labeling as peptidoglycan is actively remodeled. Yellow triangles with black outline indicate loss of TADA signal in YB642 (WT) as stalk is extended from the base. Cells were grown in rich medium (PYE) to saturation and sub-cultured into phosphate limited (HIGG) medium at 26°C for 60h before labeling. To label whole cells, 250 μM TADA was added and cells were allowed to grow an additional ~12-14h (overnight). Cells were then washed 2X with PYE to remove TADA and imaged via time-lapse with phase and fluorescence microscopy. Scale bars = 2 μm. (E) Pulse-chase time-lapse of FDAA (TADA) labeling for strains YB8597 (Δ*bacA*) showing elongation and septation of a pseudo-stalk. Yellow triangles with black outline indicate the septation site. Cells were grown as described in Figure 6D. Scale bars = 2 μm. (F) DAPI staining for DNA in strains YB642 (WT) and YB8597 (Δ*bacA*) showing DNA is present in the pseudo stalk (yellow triangles). Representative images are shown. Scale bars = 2 μm. See also Video S7.

There are several modes of PG synthesis that could result in pseudostalks. For example, they could be synthesized 1) from the base like WT stalks, albeit with a wider and variable area of extension; 2) via polar growth similar to vegetative growth of the filamentous Streptomycetes [3], with PG remodeling occurring through polar tip extension; or 3) be the result of dispersed PG synthesis throughout the pseudostalk, similar to lateral elongation of some rod-shaped cells [3].

To test these possibilities, we employed various methods of Fluorescent D-Amino Acid (FDAA) [33] labeling to visualize active PG synthesis during growth in low-phosphate medium. Cells were first subjected to short labeling pulses (45 min or ~20% of doubling time) of a red FDAA (TADA) (Figure 6B). If pseudostalks are synthesized from the base, the FDAA should label the cell and stalk-to-cell body junction, but not the pseudostalk itself. In WT cells, labeling was diffuse throughout the cell, but absent from the stalks, except from the stalk-to-cell body junction where stalk PG is synthesized (Figure 6B), consistent with previous results [11]. In contrast, Δ*bacA* cells showed strong labeling throughout the pseudostalks, indicating that these structures are not solely synthesized from the base as in WT stalks (Figure 6B). Because the labeling time was short relative to the time required for pseudostalk growth, we hypothesized that pseudostalk PG synthesis occurs through PG synthesis dispersed throughout the structure rather than by tip PG synthesis. To obtain better spatiotemporal resolution of PG remodeling and to correlate SpmX localization with PG remodeling, we performed virtual time-lapse FDAA labeling, where short successive pulses of different colored FDAA probes are used for spatiotemporal labeling of areas of PG synthesis. *A. biprosthecum* WT SpmX-eGFP and Δ*bacA* SpmX-eGFP cells were first labeled with a blue FDAA (HADA) for 6 min followed by a red FDAA (TADA) for 3 min, with the dye being washed away after each respective incubation (Figure 6C). Cells were then imaged via 3D-SIM (<u>S</u>tructured <u>I</u>llumination <u>M</u>icroscopy). For WT cells, the HADA and TADA signals overlapped both in the cell body and at the base of the stalk, colocalized with the SpmX-eGFP focus and consistent with stalk PG synthesis at the base and no PG synthesis along the stalk length (Figure 6C). In contrast, the HADA and TADA signals overlapped in both the cell body and the pseudostalks of Δ*bacA* cells and with SpmX-eGFP localizing to the tips of the pseudostalks (Figure 6C), indicating that pseudostalks are formed via dispersed PG synthesis throughout the structure.

### BacA inhibits the default elongation and division modes of PG synthesis as well as DNA entry at sites of stalk synthesis

While the FDAA labeling techniques used above (Figures 6B & 6C) show that new PG is incorporated into the pseudostalk, it does not distinguish *per se* if PG is being incorporated from the base, as in WT stalks, or if PG incorporation is distributed throughout the pseudostalk. To determine the PG incorporation pattern for Δ*bacA* pseudostalks, we performed pulse-chase FDAA labeling (Figure 6D). *A. biprosthecum* WT and Δ*bacA* cells were first labeled with a red FDAA (TADA) for ~12-14 hours (~3-3.5 cell cycles) to ensure whole-cell labeling, including stalks and pseudostalks. Cells were then washed twice to remove the FDAA and time-lapse fluorescent microscopy was performed. In WT cells, stalks are synthesized from the base [11]. Once PG is added to the stalk, it is inert; that is, stalk PG is not diluted by the insertion of new material, nor removed by recycling. As the *A. biprosthecum* WT cells grew, FDAA signal disappeared from the cell body, but was retained in the stalk (Figure 6D), with a slight clearing near the base as stalk PG was synthesized from its base (Figure 6D, yellow triangles). In contrast, Δ*bacA* cells lost FDAA signal throughout both the cell body and the pseudostalks at a similar rate (Figure 6D), consistent with PG synthesis throughout the pseudostalk (Figure 6B). Strikingly, 49% (42/85) of Δ*bacA* cells observed were able to extend and divide from the pseudostalks and produced cell-like extensions that continued to elongate after cytokinesis (Figure 6E, yellow triangles), suggesting that cell growth was occurring at sites usually reserved for stalk synthesis. These results suggested that BacA might be required to prevent entry of chromosomal DNA that would be required for continued growth of these lateral cell extensions. To determine if Δ*bacA* pseudostalks contained DNA, we stained live cells with DAPI. In WT cells, DAPI staining was constrained to the cell body, but in Δ*bacA* cells, pseudostalks stained strongly for DNA (Figure 6F). Stalks are normally compartmentalized from the cytoplasm [9], but in the absence of BacA, this appears to no longer be the case. It is unclear if BacA plays a direct role in excluding DNA from the stalk or if this is an indirect consequence of BacA organizing other stalk synthesis proteins at the base. It should be noted that we were unable to determine the viability of the cell-like structures produced from the pseudostalks, but, qualitatively, there is no discernable difference between the growth of WT and Δ*bacA* strains in liquid culture or on plates. Taken together, these data indicate that, in addition to providing a scaffold for proper stalk PG synthesis at its base, BacA inhibits not only unwanted PG synthesis involved in cell elongation and division, but also the entry of DNA at sites of stalk synthesis.

## Discussion

Intricate biological processes underlie even the simplest of shapes. The ubiquitous rod- and sphere-shaped bacteria can be generated through multiple strategies [34, 35], but the final observed form is the result of finely tuned gene expression, metabolic processes that regulate PG precursor and subunit levels, and the spatiotemporal localization of PG-modifying enzymes. The study of shape generation and maintenance has uncovered a common theme of scaffolding proteins such as MreB, FtsZ, and DivIVA that organize morphogenic, PG remodeling processes. These cytoskeletal scaffolding proteins are 1) required for the recruitment of downstream enzymes and accessory proteins involved in cell wall remodeling and 2) control the spatial activity of these proteins. It should be noted that the PG remodeling processes are not necessarily discrete and must often be regulated in relation to each other. For instance, during a large portion of the cell cycle in *C. crescentus*, FtsZ serves as the most upstream localization factor for MreB, which is responsible for elongation and maintenance of cell width [36, 37].

Here we present evidence that the paradigm of a cytoskeletal scaffolding protein serving the dual roles of recruitment and organization in PG synthesis is not always true. In *A. biprosthecum*, the roles of recruitment and organization for the stalk synthesis PG remodeling complex are performed by two distinct proteins, SpmX and BacA, respectively. We propose a model in which SpmX recruits but does not organize and BacA organizes but does not recruit. Furthermore, BacA prevents cell elongation and division associated PG synthesis from being activated at the site of stalk PG synthesis. Finally, BacA also prevents the entry of DNA into stalks.

Previous work and this study show that SpmX appears early in *Asticcacaulis* spp. cell cycle, where it localizes in pre-divisional daughter cells to mark the future site of stalk synthesis ([11] and Figures 4A & 7A). Here we show that BacA operates downstream of SpmX for subcellular localization in the *A. biprosthecum* stalk synthesis pathway but acts upstream of SpmX for the topological organization of SpmX and the PG synthesis machinery. In the absence of SpmX, cells are stalkless ([11] and Figures 3A & 4G) and BacA-mVenus forms foci that move randomly throughout the cell body, indicating that SpmX recruits BacA, and presumably other PG remodeling enzymes that remain to be discovered, to the site of stalk synthesis (Figure 7B). In a Δ*bacA* Δ*spmX* double mutant, cells phenocopy the Δ*spmX* mutant and fail to produce stalks or pseudostalks (Figure 4G). This supports the conclusion that BacA operates downstream of SpmX. The SpmX muramidase domain has both PG binding and hydrolysis activity, indicating that it is in the periplasm [15], whereas BacA is cytoplasmic, making their interaction puzzling. However, results from a number of papers indicate that the *C. crescentus* SpmX muramidase domain spends at least some time in the cytoplasm, where it can interact with BacA [30, 38–40]. We hypothesize that the same is true for *A. biprosthecum*.

**Figure 7.**
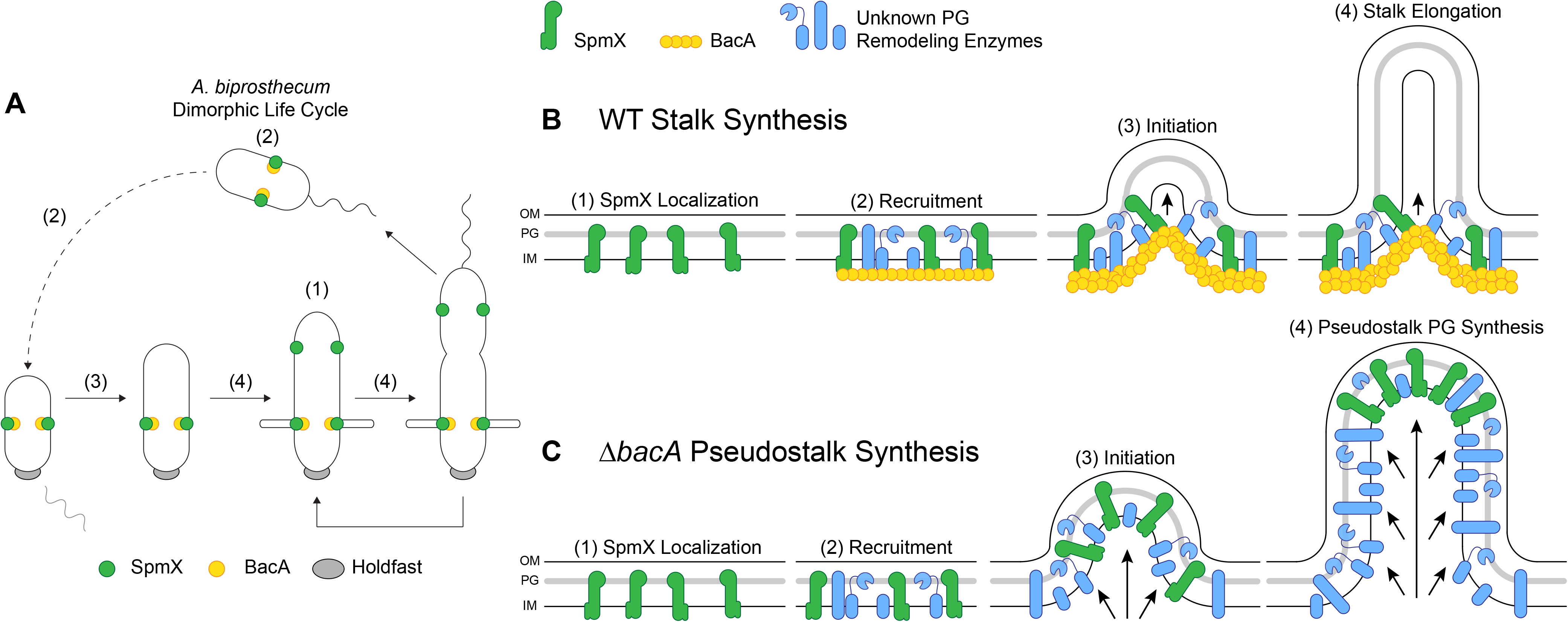
Model of the Stalk PG Biosynthetic Complex in *A. biprosthecum*. (A) Dimorphic life cycle of *A. biprosthecum*. The prosthecate mother cell produces an adhesive holdfast (shown in gray) at one pole. Cell division results in a motile, nonreplicating swarmer cell that differentiates into a prosthecate cell. Green circles indicate SpmX localization. Yellow circles represent BacA localization. Numbers in parentheses, (1)-(4), indicate stages of WT stalk synthesis as described in Figure 7B. (B) Model of *A. biprosthecum* WT stalk synthesis. (1) SpmX (green) appears early in the cell cycle, prior to cell division, where it marks the future site of stalk synthesis. (2) SpmX recruits the putative PG remodeling complex (blue) and the BacA scaffold (yellow) to the site of stalk synthesis. (3) PG synthesis is initiated at the site defined by SpmX localization and in an area constrained by BacA. (4) As PG synthesis progresses and the stalk elongates, synthesis is constrained at the base of the stalk by the BacA scaffold. Timing of these stages in the cell cycle are indicated by the matching numbers in Figure 7A. (C) Model of *A. biprosthecum* Δ*bacA* pseudostalk synthesis. As with WT cells, (1) SpmX (green) appears early in the cell cycle, prior to cell division, where it marks the future site of stalk synthesis. (2) SpmX recruits the putative PG remodeling complex (blue), but without BacA. (3) PG synthesis is initiated at the site defined by SpmX localization. (4) As PG synthesis progresses, there is no BacA scaffold to constrain the complex at the base of the synthesis site; PG synthesis occurs promiscuously throughout, with SpmX localized to the distal end of the structure, leading to short, wide pseudostalks. Timing of these stages in the cell cycle are indicated by the matching numbers in Figure 7A.

Interestingly, once stalk synthesis is initiated, BacA serves as a scaffolding protein that constrains PG synthesis at the base of the stalk, allowing for WT stalk elongation and maturation (Figure 7B). Based on current knowledge, this is the first reported morphogenic PG remodeling process in which the protein responsible for recruitment of the complex is different from the scaffolding protein that organizes the complex. The stalkless phenotype of *spmX* and *bacA* mutants in *A. biprosthecum* is striking compared to *C. crescentus*, where neither gene is required for stalk synthesis. Deletion of *C. crescentus bacA*_*Cc*_ does not impact stalk synthesis in the complex PYE medium and leads to a mild reduction of stalk length only under the strong stalk elongation stimulating conditions of phosphate starvation [12, 16]. In *A. excentricus*, which has a single sub-polar stalk, *spmX* is required for sub-polar stalk synthesis, but phosphate starvation results in polar stalk synthesis in the *spmX* mutant [11]. Perhaps the synthesis of non-polar stalks has additional topological constraints that are more easily solved by the separation of the recruitment and scaffolding processes into two different proteins as we have described here, as this separation may be more adaptable for generating unique morphologies. Comparative analysis of the requirements for stalk synthesis in these three species provides an excellent model to determine how evolution can solve different morphological constraints.

Many questions remain regarding prosthecae synthesis in both *A. biprosthecum* specifically and the stalked Alphaproteobacteria in general. Even among closely related species there are phenotypic differences, such as the number and placement of stalks, as well as differences in the genes involved in stalk synthesis. Furthermore, in many stalked marine species, the stalk serves a reproductive function, with a budding daughter cell produced from the tip of the stalk [41–44]. How have these stalk synthesis pathways evolved? Are there core genes that are shared amongst bacteria that produce stalks? There are certainly differences, specifically between *C. crescentus* and *A. biprosthecum* (Table S1) [11], but what other genes are involved in *A. biprosthecum* stalk synthesis? ABI_34190, the gene that lies upstream and overlaps *bacA* (Figure 1C), is a putative M23 family metallopeptidase (Pfam 01551) whose predicted endopeptidase activity would cleave PG crosslinks. Thus, ABI_34190 provides an intriguing candidate for both its predicted function in PG remodeling and its genomic association with *bacA*. How do bactofilins function in stalk synthesis? Bactofilin’s ability to polymerize appears to be important for function. Polymerization is thought to be mediated via hydrophobic interactions between conserved hydrophobic residues in the core DUF583 domain [28]. Point mutations in *C. crescentus* BacA identified residues that disrupt polymerization *in vitro* and are important for localization to the stalked pole [28]. Analogous mutations in the *H. pylori* bactofilin CcmA also disrupt polymerization *in vitro*, result in mislocalization *in vivo*, and, when expressed as the sole copy of CcmA, the mutant strains are morphologically indistinguishable from a *ccmA* null mutant [45]. Taken together, these results show that bactofilin polymerization can be easily disrupted by mutating selected residues and that polymerization is required for proper localization and function *in vivo*. The terminal regions flanking the conserved bactofilin domain often vary between species. Do the N- and C-terminal domains serve any specific purposes? Work on *T. thermophilus* bactofilin showed that the N-terminal region is involved in membrane binding [31]. We used high detergent levels, a high micellization condition, to get purified BacA to form filaments *in vitro*. Our BACTH results suggest that the C-terminus is important for BacA-SpmX interaction. It seems likely that these N- and C-terminal regions play important roles in membrane and protein interactions.

This study has also revealed an unexpected role for a scaffold protein in inhibiting default modes of PG synthesis. In the absence of BacA, cells produce what we have termed “pseudostalks”, which are significantly shorter and wider than stalks. In addition, BacA is required to constrain SpmX at the base of the stalk (Figure 7B), as observed by the mislocalization of SpmX-eGFP in the absence of BacA (Figure 7C). Pseudostalks, like WT stalks, are composed of PG. However, unlike WT stalks, which are synthesized from the base of the stalk (Figure 7B), pseudostalks are the result of dispersed PG remodeling throughout the structure (Figure 7C). Thus, SpmX is sufficient to recruit PG synthesis enzymes to the pseudostalk in the absence of BacA, however we do not know yet if these are the same enzymes that normally drive stalk synthesis. We find a parallel for this phenomenon with the cytoskeletal scaffolding protein DivIVA in the Actinobacterium *Mycobacterium smegmatis*. Normal, rod-shaped *M. smegmatis* undergo asymmetric bipolar growth, and DivIVA focuses this cell envelope assembly at the poles [46]. When DivIVA is depleted, PG synthesis persists, but in a disorganized manner throughout the cell, leading to spherical cells [47]. Similarly, in the absence of BacA, PG synthesis persists, but in a disorganized manner leading to pseudostalks.

Strikingly, we show that the pseudostalks that form in the absence of BacA can incorporate the default cell elongation and cell division modes, forming cell body extensions that contain DNA and can divide. These results suggest that the cytoskeletal bactofilin BacA not only organizes and constrains PG synthesis to specific subcellular locations (Figure 7B) but also inhibits PG synthesis associated with cell elongation and division and DNA entry at the site of stalk synthesis defined by SpmX. The decoupling of the recruitment and scaffolding roles for proteins that localize the PG synthesis complex may facilitate safeguarding against establishment of cell elongation and division at those morphogenetic sites. However, removal of a single layer of regulation (in this case, *A. biprosthecum* BacA) can lead to unregulated growth, with *A. biprosthecum* becoming a branching organism via pseudostalks. This follows an emerging theme of bacterial morphogenesis in which morphological variation results from the differential regulation of PG remodeling [3, 48, 49]. In other words, much like the Actinobacteria *Streptomyces* spp., *A. biprosthecum* localizes PG synthesis at specific places with different morphological results: branching hyphae and stalks, respectively. Furthermore, this result suggests that at least some of the PG synthesis enzymes used for stalk synthesis are the same as those used for cell elongation and division, which may explain why we have yet to identify a PG synthesis enzyme used specifically for stalk synthesis.

## Acknowledgements

The NIH supported this work with grants 2R01GM051986 and R35GM122556 (to Y.V.B.) and GM113172 to M.S.V. and Y.V.B. Y.V.B. is also supported by a Canada 150 Research Chair in Bacterial Cell Biology. Structured illumination microscopy (3D-SIM) data acquisition was supported by NIH grant NIH1S10OD024988-01 to the Indiana University Bloomington Light Microscopy Imaging Center. We thank Barry Stein of the Indiana University Bloomington Electron Microscopy Center for training and assistance. We wish to thank members of the Brun Lab for support, advice, and encouragement, and, in particular, Cécile Berne, Marie Delaby, Kelley Gallagher, David Kysela, Emily Sprowls, Liu Yang, and Sébastien Zappa for critical manuscript reading and feedback. We thank Merrin Joseph of the (Malcolm) Winkler Lab (Indiana University) for rotation work that contributed to strain generation for this study. Wholehearted thanks for critical preprint review by the journal club of the (Pamela) Brown Lab (University of Missouri).

## Author Contributions

Conceptualization, P.D.C. and Y.V.B.; Methodology, P.D.C., M.J., and Y.V.B.; Investigation, P.D.C. and M.J.; Resources, Y.V.B. and fluorescent D-amino acids (FDAAs) from M.S.V.; Writing – Original Draft, P.D.C; Writing – Review and Editing, P.D.C., M.J., and Y.V.B.; Visualization, P.D.C.; Supervision, Y.V.B.; Funding Acquisition, Y.V.B.

## STAR Methods

### RESOURCE AVALABILITY

#### Lead Contact

Further information and requests for resources and reagents should be directed to and will be fulfilled by the Lead Contact, Yves V. Brun.

#### Materials Availability

Unique plasmids and bacterial strains generated in this study are available upon request from the Lead Contact.

#### Data and Code Availability

Source data used in this paper for Figures 2C-2F, 3G, 4E-4F, S1, and S2 are available at Caccamo, Paul (2020), “A Division of Labor in the Recruitment and Topological Organization of a Bacterial Morphogenic Complex – Raw Data”, Mendeley Data, v1. http://dx.doi.org/10.17632/6g8zx4r6kj.1

### EXPERIMENTAL MODEL AND SUBJECT DETAILS

All freezer stocks were maintained in 10% DMSO at −80°C*. A. biprosthecum* strains used in this study were grown in liquid PYE medium at 26°C supplemented with antibiotics or supplements as necessary (kanamycin 5 μg ml^−1^, gentamicin 0.5 μg ml^−1^, spectinomycin 25 μg ml^−1^, streptomycin 5 μg ml^−1^, and 0.3 mM diaminopimelic acid). Strains were maintained on PYE plates at 26°C supplemented with antibiotics or supplements as necessary (kanamycin 5 or 20 μg ml^−1^, gentamicin 2.5 μg ml^−1^, spectinomycin 50 μg ml^−1^, streptomycin 20 μg ml^−1^, 3% sucrose, and 0.3 mM diaminopimelic acid). For phosphate starvation, cells were grown in <u>H</u>utner base-<u>i</u>midazole-buffered-<u>g</u>lucose-<u>g</u>lutamate (HIGG) medium [54] containing 30 μM phosphate (phosphate-limited), supplemented with biotin at 40 ng ml^−1^ and antibiotics where appropriate. For microscopy of PYE grown samples, strains were inoculated in PYE from colonies and grown at 26°C with shaking until mid- to late-exponential phase (OD_600_ ≥ 1.0) before imaging. For microscopy of HIGG grown, phosphate-limited samples, strains were inoculated in PYE from colonies and grown at 26°C with shaking until late-exponential phase. Cultures were then washed 2X with deionized distilled H_2_O (ddH_2_O), diluted 1:20 in HIGG, and grown at 26°C with shaking for 72h before imaging. *E. coli* strains used in this study grown in liquid <u>l</u>ysogeny <u>b</u>roth (LB) medium at 37°C (or 30°C for BACTH assays) supplemented with antibiotics or supplements as necessary (ampicillin 100 μg ml^−1^, kanamycin 30 μg ml^−1^, gentamicin 15 μg ml^−1^, spectinomycin 100 μg ml^−1^, streptomycin 30 μg ml^−1^, 0.3 mM diaminopimelic acid, and X-gal 40 μg ml^−1^). Strains were maintained on LB plates at 37°C (or 30°C for BACTH assays) supplemented with antibiotics or supplements as necessary (ampicillin 100 μg ml^−1^, kanamycin 25 or 50 μg ml^−1^, gentamicin 20 μg ml^−1^, streptomycin 30 or 100 μg ml^−1^, 0.3 mM <u>d</u>i<u>a</u>mino<u>p</u>imelic acid (DAP), X-gal 40 μg ml^−1^, and 0.5 mM IPTG). Electroporation of *A. biprosthecum* was performed as previously described [55]. Outgrowth was performed for 8-24h at 26°C. For electroporation with replicating plasmids, 100 μl of outgrowth culture was plated on PYE plates with appropriate selection at dilutions of 10^1^, 10^0^, and 10^−1^. For electroporation with integrating plasmids, outgrowth was divided into volumes of 100, 300, and 600 μl and plated on PYE plates with appropriate selection. In-house stocks of chemically competent BL21lDE3 (YB1000), DAP auxotroph WM3064 (YB7351), XL-1 Blue (YB0041), and BTH101 (YB9171) *E. coli* cells were prepared as previously described [56, 57]. A detailed list of strains is included as **Table S3**.

### METHOD DETAILS

#### Recombinant DNA Methods

DNA amplification, Gibson cloning, and restriction digests were performed according to the manufacturer. Restriction enzymes and Gibson cloning mix were from New England Biolabs. Cloning steps were carried out in *E. coli* (alpha-select competent cells, Bioline or XL1-Blue, Stratagene) and plasmids were purified using Zyppy Plasmid Kits (Zymo Research Corporation). Sequencing was performed by the Indiana Molecular Biology Institute and Eurofins MWG Operon Technologies with double stranded plasmid or PCR templates, which were purified with a DNA Clean & Concentrator kits (Zymo Research Corporation). Chromosomal DNA was purified using the Bactozol Bacterial DNA Isolation Kit (Molecular Research Center). Plasmids were introduced into all *E. coli* strains using chemical transformation according to the manufacturer’s protocols. Electroporation of *A. biprosthecum* was performed as previously described [55]. Outgrowth was performed for 8-24h at 26°C. For electroporation with replicating plasmids, 100 μl of outgrowth culture was plated on PYE plates with appropriate selection at dilutions of 10^1^, 10^0^, and 10^−1^. For electroporation with integrating plasmids, outgrowth was divided into volumes of 100, 300, and 600 μl and plated on PYE plates with appropriate selection. Mating of plasmids into *A. biprosthecum* was performed using the *dap*‾ *E. coli* strain WM3064 (YB7351) [58]. Allelic exchange and deletions were achieved using a two-step sucrose counterselection procedure with pNPTS138/pNPTS139. Insertional fusions of eGFP and mVenus were made using pGFPC-1 and pVENC-4, respectively [59]. BACTH plasmids were made using the Euromedex BACTH System Kit (Euromedex Cat. No. EUK001).

#### Plasmid Construction

All PCR, including insert fragments for Gibson assembly, was performed using iProof High-Fidelity DNA Polymerase (Bio-Rad Laboratories, Inc., Cat. No. 1725302) according to the manufacturer’s instructions. All plasmids were constructed via Gibson assembly using NEBuilder HiFi DNA Assembly Master Mix (New England Biolabs, Inc., Cat. No. E2621X) according to the manufacturer’s instructions. Prior to use in cloning, all restriction enzyme digested plasmids were treated with Calf Intestinal Alkaline Phosphatase (New England Biolabs, Inc., Cat. No. M0290S) according to the manufacturer’s instructions. All primers for Gibson assembly were designed using NEBuilder Assembly Tool (New England Biolabs, Inc.). For the *bacA* complementation vector, we fused the promoter region, comprising 236 bp immediately upstream of ABI_34190, to the coding region of *bacA* and placed the promoter/gene fusion on the low copy plasmid pMR10. The promoter/gene fusion is placed in the opposite orientation of the lacZ promoter. The restriction enzymes used are as follows for pNPTS138/pNPTS139-based vectors (*Eco*RV or *Sph*I/*Nhe*I), pET28a+ (*Sal*I/*Sac*I), pMR10 (*Eco*RV), pVENC-4 *(Nde*I/*Kpn*I), and BACTH plasmids (*Kpn*I/*Eco*RI). A detailed list of plasmids and primers can be found in the **STAR Methods Key Resources Table**, **Table S2 (plasmids)**, and **Table S4 (primers)**.

#### Microscopy

##### Light Microscopy and Fluorescence Imaging

For light microscopy analysis, cells were spotted onto pads made of 1% SeaKem LE Agarose (Lonza, Cat. No. 50000) in PYE and topped with a glass coverslip. When appropriate, the coverslip was sealed with VALAP (<u>va</u>seline, <u>la</u>nolin, and <u>p</u>araffin at a 1:1:1 ratio). Images were recorded with inverted Nikon Ti-E microscopes using either 1) a Plan Apo 60X 1.40 NA oil Ph3 DM objective with DAPI/FITC/Cy3/Cy5 or CFP/YFP/mCherry filter cubes and an iXon X3 DU885 EMCCD camera or 2) a Plan Apo λ 100X 1.45 NA oil Ph3 DM objective with DAPI/FITC/Cy3/Cy5 or CFP/YFP/mCherry filter cubes and a Photometrics Prime 95B sCMOS camera. Images were processed with NIS Elements software (Nikon). To visualize DNA, cells were grown in HIGG medium for 72h as described above. Cells were washed 1x with PYE and 1 μl of cell suspension was spotted onto a coverslip and topped with an agar pad. Cells were monitored via phase microscopy for ~1h to ensure they were growing and 20 μl of 1 mg/ml of 4’,6-diamidino-2-phenylindole dihydrochloride (DAPI) was spotted on top of the agar pad. DAPI was allowed to diffuse through the pad to the cells and cells were imaged.

##### Structured Illumination Microscopy

3D-SIM images were collected on a DeltaVision OMX system (Applied Precision Inc., Issaquah, USA) equipped with a 1.4NA Olympus 100X oil objective. A 405 nm laser source and 419-465 nm emission filter were used for collecting HADA signal. A 488 nm laser source and 500-550 nm emission filter were used for collecting GFP signal. A 568 nm laser source and 609-654 emission filter were used for collecting TADA signal. The z-axis scanning depth was 2 μm. Immersion oil with refraction index of 1.514 was used. SIM images were reconstructed using softWoRx.

##### Electron Microscopy

10 μl cell or sacculi suspension was applied to an electron microscopy grid (Formvar/Carbon on 300 mesh; Ted Pella Inc., Cat. No. 01753-F) for 5 min at room temperature. Excess liquid was removed with Whatman filter paper. Cells or sacculi were then negatively stained with 10 μl 1% uranyl acetate (UA) and excess UA liquid was immediately removed with Whatman filter paper. Grids were allowed to dry, stored in a grid holder in a desiccation chamber, and imaged with a kV JEOL JEM 1010 transmission electron microscope (JEOL USA Inc.). Protein visualization was performed as sacculi suspension except protein was applied to the grid for 1 min and stained with 2% UA before imaging.

#### Sacculi purification

Strains were inoculated in 3 ml PYE from colonies and grown at 26°C with shaking until late-exponential phase. Cultures were then washed 2X with ddH_2_O, diluted 1:50 in HIGG, and grown at 26°C with shaking for 72h. Cells were harvested by centrifugation (7,000 g, 4°C, 15 min) and resuspended in 10 ml ddH_2_O. Cell suspension was added dropwise to 20 ml boiling 7.5% SDS in a 125 ml flask with a stir bar. Once cells addition was complete, the SDS cell suspension was boiled for 30 min with stirring and then allowed to cool to room temperature. Suspension was then washed 6X with ddH_2_O using centrifugation (100,000 g, 25°C, 30 min) to isolate the sacculi pellet and resuspended in 5 ml ddH_2_O before imaging.

#### Protein purification

Recombinant plasmids overproducing His_6_-BacA protein were transformed into BL21 *E. coli* strain. Transformants were grown, in LB medium supplemented with kanamycin (50 μg ml^−1^), at 37°C until the OD_600_ = 0.6. Expression was induced by adding 0.5 mM IPTG (isopropyl β-D-thiogalactopyranoside) and incubation was continued either 3h at 30°C or overnight at 20°C. Cells pellets were resuspended in 1/25 volume of Buffer A (TrisHCl at pH 8, 300 mM NaCl, 2 mM β-mercaptoethanol) and lysed by sonication (20 sec On / 40 sec Off, 5 min, Misonix S4000). Cells debris were pelleted by centrifugation at 36,000 g for 30 min at 4°C. The supernatant was loaded on a Ni-NTA resin (Qiagen) on an AKTA FPLC pure system. After washing with Buffer A, the protein was eluted with a linear gradient of Buffer B (50 mM TrisHCl at pH 8, 300 mM NaCl, 2 mM β-mercaptoethanol, 500 mM imidazole). Elution fraction was loaded on SDS page gel and peak fractions containing the protein were pooled. Upon purification, His_6_-BacA protein was used for polymerization assay. Polymerization assay were performed by dialyzing His_6_-BacA in Buffer C (25 mM TrisHCl at pH 8, 250 mM NaCl, 2 mM β-mercaptoethanol) supplemented with 10% Triton X-100 in the case of BacA.

#### Western blots

All strains were grown until OD=1.0 before being centrifuged and resuspended in 100 μl of 1xPBS supplemented with 0.1 μl of Universal Nuclease (Pierce #88700) sonicated for 3 times for 10sec at 50% intensity (Misonix S4000). Protein concentration was measured and normalized if needed before addition of laemmli buffer. 15 μl of each sample was loaded onto 4-20% precast polyacrylamide gels (BioRad) before being transferred on a nitrocellulose membrane according to manufacturer’s instructions. Loading was controlled by Ponceau’s staining before immunoblotting was performed by addition of an anti-GFP polyclonal antibody (MBL #598) as the primary antibody and a goat anti-rabbit HRP (Pierce) as secondary antibody. Transferred blots were visualized with SuperSignal West Pico substrate (ThermoFisher Scientific) using a Bio-Rad Chemidoc.

### QUANTIFICATION AND STATISTICAL ANALYSIS

#### Bioinformatics

Genomic and protein sequence data were obtained from the Integrated Microbial Genomes (IMG) database [60]. Multiple sequence alignment of bactofilin sequences was performed using Jalview [61] with a MAFFT alignment using the L-INS-i preset. The *C. crescentus* BacB_Cc_ protein sequence used for alignment is a translation of an open reading frame in CC3022 that initiates at a downstream ATG codon, as described previously [12]. The predicted structure of *A. biprosthecum* BacA (ABI_34180) was modeled using SWISS-MODEL [50] with the structure of *C. crescentus* BacA (CC1872; PBD code 2N3D) as a template. Alignment was then processed using the ESPript server [62]. Percent identity (PID) was calculated from pairwise MAFFT alignments using the L-INS-i preset. 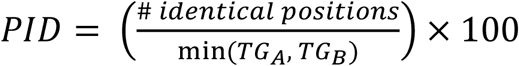, where *TG*_*A*_ and *TG*_*B*_ are the sum of the number of residues and internal gap positions in sequences A and B in the alignment [63].

#### Image analysis

Stalk length and stalk percentage data was obtained using FIJI (<u>F</u>iji <u>I</u>s <u>J</u>ust <u>I</u>mageJ) [64]. Briefly, phase micrographs were imported into FIJI and stalks were manually traced using the “Freehand Line” tool. Stalks lengths were determined from the manual trace using the “Measure” function, calibrated to the μm/pixel scale of the original micrograph. The percentage of stalked cells per image was calculated by manually counting the number of stalked and non-stalked cells per image. Subcellular localization of SpmX-eGFP and BacA-mVenus foci and subsequent localization heatmaps were generated using the ImageJ plugin MicrobeJ [65]. SpmX-eGFP maxima area and intensity were determined using the ImageJ plugin MicrobeJ [65]. Area was first generated as pixel area and converted to nm^2^ based on the pixel to μm ratio for the images. Intensity is defined as the average gray value measured on the channel used to detect the particle. Analysis of BacA-mVenus time-lapse videos and BacA-mVenus foci dynamics was performed using the ImageJ plugin TrackMate [66]. Colocalization analysis was performed by using MicrobeJ [65] to detect all cells that contained both BacA-mVenus and SpmX-mCherry foci and then measure colocalization by Pearson correlation coefficient. All statistical analysis and data visualization was generated in R version 3.5 [67] and using the ggplot2 [68] and ggsignif [69] packages. Cell silhouettes and cell body outlines were produced by importing the phase micrograph into Adobe Illustrator CC 2015.2.1 (Adobe Inc.) and manually tracing the silhouette or outline. The statistical details describing the quantification of cell morphology and fluorescent image analysis can be found in the respective figure legends.

## Supplemental Video Titles and Legends

**Video S1. Timelapse of BacA-mVenus localization WT *A. biprosthecum* cells.** Related to Figure 3 and Video S2. Subcellular localization timelapse of BacA-mVenus in strain YB7474 (*bacA*::*bacA-mVenus*) grown in rich medium (PYE) at 26°C for 24-48h. Video S1 was used to create the panels shown in Figure 3A.

**Video S2. Timelapse of BacA-mVenus localization WT *A. biprosthecum* cells.** Related to Figure 3 and Video S1. Subcellular localization timelapse of BacA-mVenus in strain YB7474 (*bacA*::*bacA-mVenus*) grown in rich medium (PYE) at 26°C for 24-48h. Video S2 is an additional timelapse of a stalked cell, showing that BacA-mVenus is functional.

**Video S3. Timelapse of BacA-mVenus localization Δ*spmX A. biprosthecum* cells.** Related to Figure 3 and Video S4. Subcellular localization timelapse of BacA-mVenus in strain YB7487 (Δ*spmX bacA*::*bacA-mVenus*) grown in rich medium (PYE) at 26°C for 24-48h. Video S3 was used to create the panels shown in Figure 3B.

**Video S4. Timelapse of BacA-mVenus localization Δ*spmX A. biprosthecum* cells.** Related to Figure 3 and Video S3. Subcellular localization timelapse of BacA-mVenus in strain YB7487 (Δ*spmX bacA*::*bacA-mVenus*) grown in rich medium (PYE) at 26°C for 24-48h. Video S4 is an additional timelapse showing that BacA-mVenus foci dynamics in the Δ*spmX* background.

**Videos S5. Timelapse of SpmX-eGFP localization in WT *A. biprosthecum* cells.** Related to Figure 4. Subcellular localization of SpmX-eGFP in strain YB5692 (*spmX*::*spmX-eGFP*). Cells were grown in rich medium (PYE) to saturation and sub-cultured into phosphate limited (HIGG) medium at 26°C for 72h. Video S5 was used to create the panels shown in Figure 4A.

**Videos S6. Timelapse of SpmX-eGFP localization in Δ*bacA A. biprosthecum* cells.** Related to Figure 4. Subcellular localization of SpmX-eGFP in strain YB7561 (*spmX*::*spmX-eGFP* Δ*bacA*). Cells were grown in rich medium (PYE) to saturation and sub-cultured into phosphate limited (HIGG) medium at 26°C for 72h. Video S6 was used to create the panels shown in Figure 4B.

**Videos S7. Timelapse of FDAA pulse chase in Δ*bacA A. biprosthecum* cells.** Related to Figure 6. Pulse-chase time-lapse of FDAA (TADA) labeling for strains YB8597 (Δ*bacA*) showing elongation and septation of a pseudo-stalk. Cells were grown in rich medium (PYE) to saturation and sub-cultured into phosphate limited (HIGG) medium at 26°C for 60h before labeling. To label whole cells, 250 μM TADA was added and cells were allowed to grow an additional ~12-14h (overnight). Cells were then washed 2X with PYE to remove TADA and imaged via time-lapse with phase and fluorescence microscopy. Video S7 was used to create the panels shown in Figure 6E.

## KEY RESOURCES TABLE

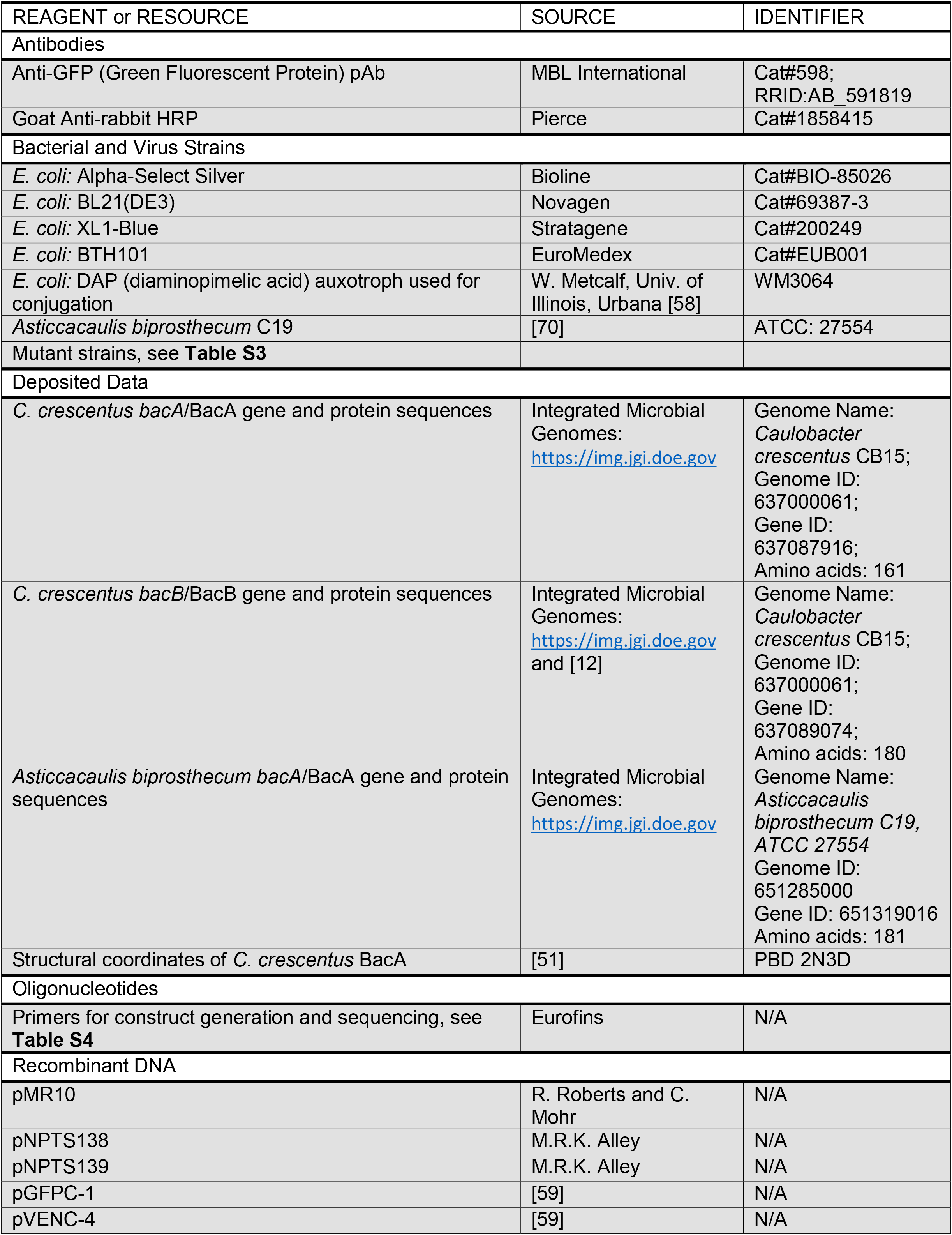
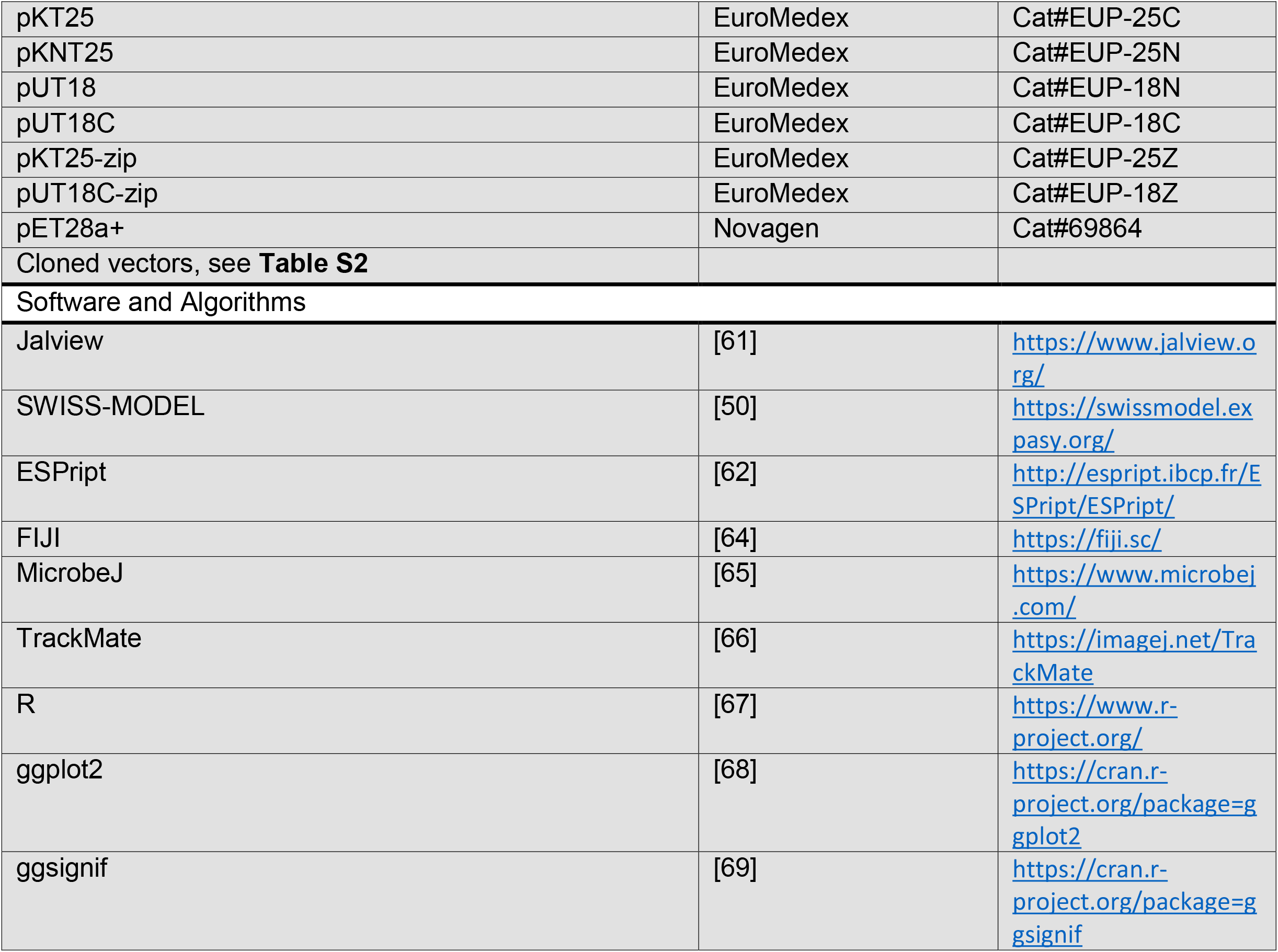

