## Supplemental Data for "A Division of Labor in the Recruitment and Topological Organization of a Bacterial Morphogenic Complex"

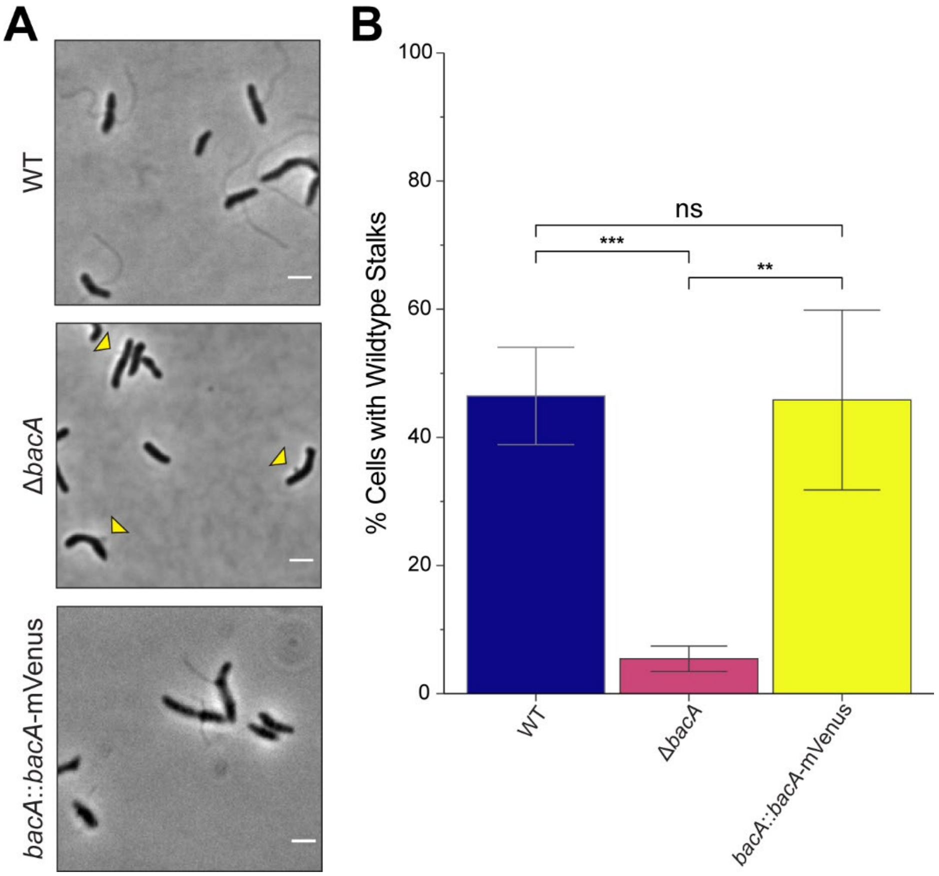


**Figure S1.** ***A. biprosthecum*** **stalk phenotypes in rich medium (PYE). Related to Figures 2 & 3.**

Analysis of stalk phenotype in strains YB642 (WT), YB8597 (Δ*bacA*), and YB7474 (*bacA*::*bacA*-mVenus). Cells were grown in rich medium (PYE) at 26°C for 48h. (A) Phase microscopy for strains YB642 (WT), YB8597 (Δ*bacA*), and YB7474 (*bacA*::*bacA*-mVenus). Scale bars = 2 µm. (B) Percentage of cells with WT stalks. Cells in phase images from Figure S1A were scored as having a WT stalk (i.e. a thin extension from the cell body). Yellow triangles indicate Δ*bacA* midcell protrusions. Data (total cells counted: WT n=747; Δ*bacA* n=577; *bacA*::*bacA*-mVenus n=378) are from four independent biological replicates with five phase microscopy fields per replicate. Data are represented as the mean (SD) percentage of cells with stalks. (***$p \leq 0.001$, **$p \leq 0.01$, ns = not significant > 0.05; two-sided t-test)

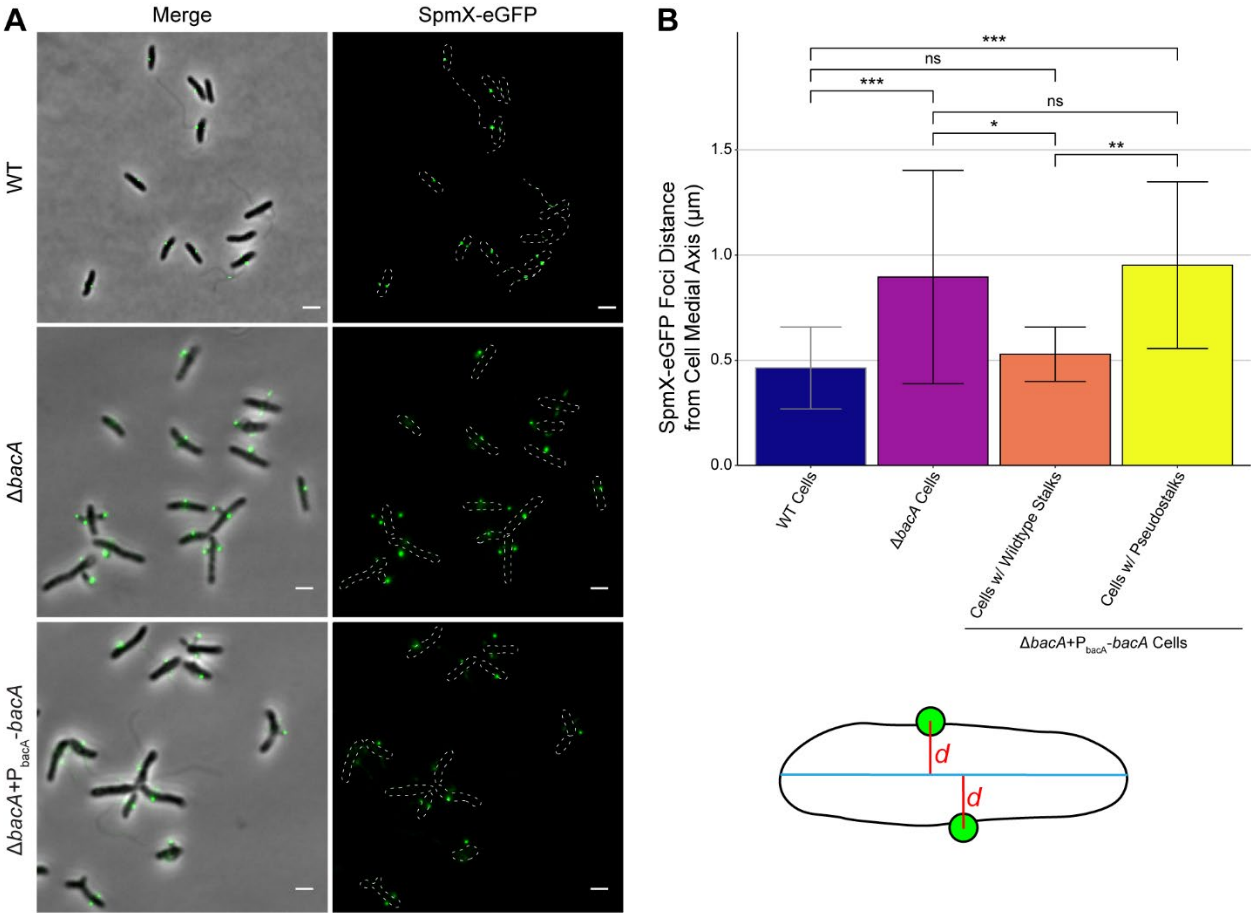


**Figure S2. SpmX Localization is Rescued for Wildtype Stalk Synthesis when BacA Is Expressed in a Δ*bacA* background. Related to Figure 4.**

(A) Subcellular localization of SpmX-eGFP in starins YB5692 (*spmX*::*spmX-eGFP*), YB7561 (*spmX*::*spmX-eGFP* Δ*bacA*), and YB9521 (*spmX*::*spmX-eGFP* Δ*bacA* + pMR10-$P_{bacA}$-*bacA*) grown in rich medium (PYE) to saturation and sub-cultured into phosphate limited (HIGG) medium at 26°C for 72h. Phase merge and fluorescence microscopy images. Scale bars = 2 μm. (B) Distribution of orthogonal distance (μm) of each SpmX-eGFP maxima from the medial axis of its associated parent cell. In order to distinguish cells with wildtype stalks from cells with pseudostalks, analysis was performed on a subset of the data presented in Figure 4C-4F. Data (WT n=49; Δ*bacA* n=22; Δ*bacA*+$P_{bacA}$-*bacA* n= 15) are plotted on a continuous y-axis and are represented as box and whisker plots as described in Figure 2D. (***$p \leq 0.001$, **$p \leq 0.01$, *$p \leq 0.05$, ns = not significant > 0.05; two-sided t-test).

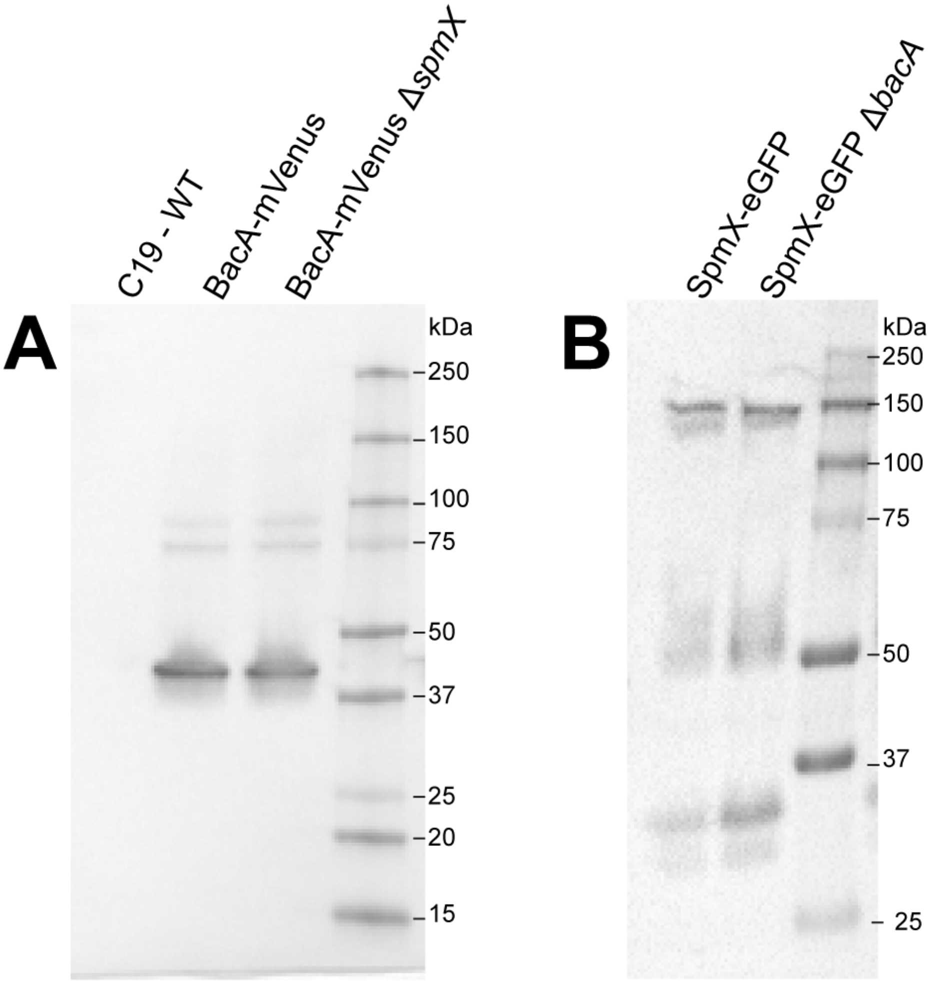


**Figure S3. Fluorescent Proteins are Expressed at Similar Levels in WT Cells and Deletion Strains. Related to Figures 3 & 4.**

(A) Western blot against BacA-mVenus inserted at the *bacA* locus in WT (YB7474) and Δ*spmX* (YB7487) cells. (B) Western blot against SpmX-eGFP inserted at the *spmX* locus in WT (YB5692) and Δ*bacA* (YB7561) cells.

| Protein | *C. crescentus* | | *A. biprosthecum* | |
|---|---|---|---|---|
| | Role in stalk synthesis | Loss of function phenotype | Role in stalk synthesis | Loss of function phenotype |
| **BacA [S1]** | • Cytoskeletal protein<br>• Localizes PbpC to stalked pole | • Shorter stalks under phosphate starvation<br>• PbpC does not localize | Unknown at time of study initiation | Unknown at time of study initiation |
| **BacB [S1]** | • Cytoskeletal protein<br>• Localizes PbpC to stalked pole | • Shorter stalks under phosphate starvation<br>• PbpC does not localize | No homolog | N/A |
| **PbpC [S1, S2]** | • Class A (bifunctional) PBP<br>• Localizes StpX to stalked pole | • Shorter stalks under phosphate starvation<br>• StpX does not localize | No homolog | N/A |
| **StpX [S2, S3]** | • Potential role in metal homeostasis | • Shorter stalks under phosphate starvation | No homolog | N/A |
| **SpmX [S4, S5]** | • None (involved in regulation of asymmetric cell division) | • Division defects | • Required for stalk synthesis<br>• Marks site of stalk synthesis | • Stalkless cells |

**Table S1. Proteins involved in stalk synthesis. Related to Figure 1.**

At the time this study was initiated, four *C. crescentus proteins* (BacA, BacB, PbpC, and StpX) had been shown to be involved in stalk synthesis. Of these four, only one (BacA) had a bi-directional best hit homolog in *A. biprosthecum*. The one *A. biprosthecum* protein involved in stalk synthesis (SpmX) has a *C. crescentus* homolog, but in *C. crescentus* this protein is involved in cell cycle regulation and not stalk synthesis.

| Plasmid | Genotype/description | Reference/source |
|---|---|---|
| pGFPC-1-spmX$_{AB}$ | pGFPC-1 bearing the C-terminal fragment of *spmX* (ABI_31540); Spec/Strep$^R$ | [S4] |
| pCHYC-1-spmX$_{AB}$ | pCHYC-1 bearing the C-terminal fragment of *spmX* (ABI_31540); Spec/Strep$^R$ | [S4] |
| pPC10 | pNPTS139 derivative used to generate a clean deletion of *bacA* (ABI_34180); Kan$^R$ | this study |
| pPC21 | pVENC-4 bearing the C-terminal fragment *bacA* (ABI_34180) $_{(nt\ 44\text{-}543)}$; Gent$^R$ | this study |
| pPC36 | Δ*bacA* complementation vector; pMR10 containing the promotor region upstream of the ABI_34190/ABI_34180 loci fused to the coding region of ABI_34180; Kan$^R$ | this study |
| pPC62 | pKT25 derivative in which SpmX is genetically fused in-frame to the T25 fragment; Kan$^R$ | this study |
| pPC63 | pKT25 derivative in which BacA is genetically fused in-frame to the T25 fragment; Kan$^R$ | this study |
| pPC65 | pKNT25 derivative in which BacA is genetically fused in-frame to the T25 fragment; Kan$^R$ | this study |
| pPC67 | pUT18 derivative in which SpmX is genetically fused in-frame to the T18 fragment; Amp$^R$ | this study |
| pPC68 | pUT18 derivative in which BacA is genetically fused in-frame to the T18 fragment; Amp$^R$ | this study |
| pPC70 | pUT18C derivative in which SpmX is genetically fused in-frame to the T18 fragment; Amp$^R$ | this study |
| pPC72 | pKNT25 derivative in which SpmX is genetically fused in-frame to the T25 fragment; Kan$^R$ | this study |
| pPC73 | pUT18C derivative in which BacA is genetically fused in-frame to the T18 fragment; Amp$^R$ | this study |
| pMJ2 | pET28a+ derivative to overexpress His-tagged BacA$_{Cc}$ (CC_1873) in *E. coli*; Kan$^R$ | this study |
| pMJ4 | pET28a+ derivative to overexpress His-tagged BacA (ABI_34180) in *E. coli*; Kan$^R$ | this study |
| YB8246 | pNPTS138-Δ*spmX*$_{Ab;}$ Kan$^R$ | [S6] |

**Table S2. Plasmids used in this study. Related to STAR Methods Key Resource Table.**

| Strain | Genotype/description | Construction | Reference/source |
| --- | --- | --- | --- |
| ***E. coli*** | | | |
| YB5681 | DH5α strain for maintaining pGFPC-1-spmX$_{AB}$ | - | [S4] |
| YB7351 | thrB1004 pro thi rpsL hsdS lacZΔM15 RP4-1360 Δ(araBAD)567 ΔdapA1341::[erm pir]; DAP (diaminopimelic acid) auxotroph used for conjugation, referred to as strain WM3064 in reference | - | [S7] |
| YB7552 | DH5α strain for maintaining pPC10 | Transformation of pPC10 into DH5α | this study |
| YB8159 | DAP auxotroph conjugative donor *E. coli* for mating plasmids into *A. biprosthecum* | Transformation of pPC10 into YB7351 | this study |
| YB8177 | DH5α strain for maintaining pPC21 | Transformation of pPC21 into DH5α | this study |
| YB8537 | DH5α strain for maintaining pMJ2 | Transformation of pMJ2 into DH5α | this study |
| YB8539 | DH5α strain for maintaining pMJ4 | Transformation of pMJ4 into DH5α | this study |
| YB8246 | DH5α strain for maintaining pNPTS138-*ΔspmX$_{Ab}$* | Transformation of pNPTS138-*ΔspmX$_{Ab}$* into DH5α | [S6] |
| YB8552 | BL21IDE3 strain containing pMJ2. For overexpression of BacA$_{Cc}$. | Transformation of pMJ2 into YB1000 | this study |
| YB8554 | BL21IDE3 strain containing pMJ4. For overexpression of BacA. | Transformation of pMJ4 into YB1000 | this study |
| YB8584 | DH5α strain for maintaining pPC36 | Transformation of pPC36 into DH5α | this study |
| YB9158 | XL1-Blue strain for maintaining pPC62 | Transformation of pPC62 into YB0041 | this study |
| YB9159 | XL1-Blue strain for maintaining pPC63 | Transformation of pPC63 into YB0041 | this study |
| YB9161 | XL1-Blue strain for maintaining pPC72 | Transformation of pPC72 into YB0041 | this study |
| YB9162 | XL1-Blue strain for maintaining pPC65 | Transformation of pPC65 into YB0041 | this study |
| YB9164 | XL1-Blue strain for maintaining pPC67 | Transformation of pPC67 into YB0041 | this study |
| YB9165 | XL1-Blue strain for maintaining pPC68 | Transformation of pPC68 into YB0041 | this study |
| YB9167 | XL1-Blue strain for maintaining pPC70 | Transformation of pPC70 into YB0041 | this study |
| YB9168 | XL1-Blue strain for maintaining pPC73 | Transformation of pPC73 into YB0041 | this study |
| ***A. biprosthecum*** | | | |
| YB5692 | C19 *spmX*::*spmX-eGFP* (Spec$^R$/Strep$^R$) | Electroporation of pGFPC-1-spmX$_{AB}$ into YB642 | [S4] |

| | | | |
|---|---|---|---|
| YB7474 | C19 *bacA*::*bacA-mVenus* (Gent$^R$) | Electroporation of pPC21 into YB642 | this study |
| YB7487 | C19 Δ*spmX* *bacA*::*bacA-mVenus* (Gent$^R$) | Electroporation of pPC21 into YB8237 | this study |
| YB7489 | C19 Δ*bacA* Δ*spmX* | Mating of YB8237 with YB8159. | this study |
| YB7561 | C19 *spmX*::*spmX-eGFP* Δ*bacA* (Spec$^R$/Strep$^R$) | Electroporation of pGFPC-1-spmX$_{AB}$ intoYB8597 | this study |
| YB8237 | C19 Δ*spmX* | Electroporation of plasmid from YB8246 into YB642 | [S6] |
| YB8597 | C19 Δ*bacA* | Mating of YB642 with YB8159 | this study |
| YB8601 | C19 Δ*bacA* / pPC36 (Kan$^R$) | Electroporation of pPC36 into YB8597 | this study |
| YB8620 | C19 Δ*bacA* / pMR10 (Kan$^R$) | Electroporation of pMR10 into YB8597 | this study |
| YB9183 | C19 / pMR10 (Kan$^R$) | Electroporation of pMR10 into YB642 | this study |
| YB9466 | C19 *bacA*::*bacA-mVenus* *spmX*::*spmX-mCherry* (Spec$^R$/Strep$^R$) | Electroporation of pCHYC-1-SpmX$_{AB}$ plasmid into YB7474 | this study |
| YB9521 | C19 *spmX*::*spmX-eGFP* Δ*bacA* (Spec$^R$/Strep$^R$) / pPC36 (Kan$^R$) | Electroporation of pPC36 into YB7561 | this study |

**Table S3. Strains used in this study. Related to STAR Methods Key Resource Table.**

| Primer ID | Sequence (5' to 3')[1] | Restriction site | Purpose |
|---|---|---|---|
| ABI_34180SeqUp | CAGGCGTTGTCTCCTTTGTC | - | Sequencing the ABI_34180 (*bacA*) locus |
| ABI_34180SeqDn | CGACTGGACGGTTGAAAAGG | - | |
| ABI_34190SeqUp | GTTAAACCTTGCGCCGTTCG | - | Sequencing the ABI_34190 locus |
| ABI_34190SeqDn | GGCACTGCGGTTGAGATTGA | - | |
| AbiBacA500C-V4F | gagacgtccaattg**catatg**AACCGACCCCGGCTCCGC | NdeI | Generation of pPC21 |
| AbiBacA500C-V4R | tcgagatcttaa**ggtacc**CTTCAGGTCGCCCAGAGCG | KpnI | |
| ABI_PlmdC-bacA_F | acgccaagcttccatgggatCTACTTCAGGTCGCCCAG | - | Generation of pPC36 |
| AbiPlytMbacA2F | CGGCGGTTATGTTCAACAAAACTAACAAACCGG | - | |
| AbiPlytMbacA1R | GTTGAACATAACCGCCGCCACCTAAAC | - | |
| ABI_PlmdC_R | gctctgcaggagatctcgatAATGTCCTTATGCAATCC | - | |
| delABI34180upF | aattctggatccacgatTCAATGTCGACGAAGACTTC | - | Generation of pPC10 |
| delABI34180upR | GAGGTGGATTAGTTTTGTTGAACATAATCTCCTG | - | |
| delABI34180dnF | ACAAAACTAATCCACCTCACCGAACTTAATCTGTGAAGAAAGC | - | |
| delABI34180dnR | aagcttcctgcaggatGCGGATTCGACCGTGCCG | - | |
| oMJ3_BacAcc | cga**gagctc**ATGTTCAGCAAGCAAGCTAAATCG | SacI | Generation of pMJ2 |
| oMJ4_BacAcc | taa**gtcgac**TTAGCCGGCGCTCTTGG | SalI | |
| oMJ5_BacAab | aga**gagctc**ATGTTCAACAAAACTAACAAACCGG | SacI | Generation of pMJ4 |
| oMJ6_BacAab | ata**gtcgac**CTACTTCAGGTCGCCCAG | SalI | |
| pUT18C_AbiSpmX-F | tctagaggatccccg**ggtacc**gGTGATGAGAGCGCGTCAAAAG | KpnI | Generation of pPC62 |
| pKT25_AbiSpmX-R | acgttgtaaaacgacggcc**gaattc**TCAGTCCTTGAGGCCGCC | EcoRI | |
| pKT25_AbiBacA-F | tctagaggatccccg**ggtacc**tATGTTCAACAAAACTAACAAACC | KpnI | Generation of pPC63 |
| pKT25_AbiBacA-R | acgttgtaaaacgacggcc**gaattc**CTACTTCAGGTCGCCCAG | EcoRI | |
| pKTN25_AbiBacA-F | tctagaggatccccg**ggtacc**gATGTTCAACAAAACTAACAAACCGG | KpnI | Generation of pPC65 |
| pKTN25_AbiBacA-R | cgattgctgcatggtcatt**gaattc**gaCTTCAGGTCGCCCAGAGC | EcoRI | |

| | | | |
|---|---|---|---|
| pUT18C_AbiSpmX-F | tctagaggatccccg**ggtacc**gGTGATGAGAGCGCGTCAAAAG | KpnI | Generation of pPC67 |
| pUT18_AbiSpmX-R | cgtggcctcgctggcggct**gaattc**gaGTCCTTGAGGCCGCCAAG | EcoRI | |
| pKTN25_AbiBacA-F | tctagaggatccccg**ggtacc**gATGTTCAACAAAACTAACAAACCGG | KpnI | Generation of pPC68 |
| pUT18_AbiBacA-R | cgtggcctcgctggcggct**gaattc**gaCTTCAGGTCGCCCAGAGC | EcoRI | |
| pUT18C_AbiSpmX-F | tctagaggatccccg**ggtacc**gGTGATGAGAGCGCGTCAAAAG | KpnI | Generation of pPC70 |
| pUT18C_AbiSpmX-R | attacttagttatatcgat**gaattc**gaTCAGTCCTTGAGGCCGCC | EcoRI | |
| pUT18C_AbiSpmX-F | tctagaggatccccg**ggtacc**gGTGATGAGAGCGCGTCAAAAG | KpnI | Generation of pPC72 |
| pKTN25_AbiSpmX-R | cgattgctgcatggtcatt**gaattc**gaGTCCTTGAGGCCGCCAAG | EcoRI | |
| pKTN25_AbiBacA-F | tctagaggatccccg**ggtacc**gATGTTCAACAAAACTAACAAACCGG | KpnI | Generation of pPC73 |
| pUT18C_AbiBacA-R | attacttagttatatcgat**gaattc**gaCTACTTCAGGTCGCCCAG | | |

**Table S4 – Primers used in this study. Related to STAR Methods Key Resource Table.**

[1]Restriction sites in **bold**; Genomic sequence in CAPS

<u>**Supplemental References**</u>


S1. Kuhn, J., Briegel, A., Morschel, E., Kahnt, J., Leser, K., Wick, S., Jensen, G.J., and Thanbichler, M. (2010). Bactofilins, a ubiquitous class of cytoskeletal proteins mediating polar localization of a cell wall synthase in *Caulobacter crescentus*. EMBO J *29*, 327-339.

S2. Hughes, H.V., Lisher, J.P., Hardy, G.G., Kysela, D.T., Arnold, R.J., Giedroc, D.P., and Brun, Y.V. (2013). Co-ordinate synthesis and protein localization in a bacterial organelle by the action of a penicillin-binding-protein. Mol Microbiol *90*, 1162-1177.

S3. Hughes, H.V., Huitema, E., Pritchard, S., Keiler, K.C., Brun, Y.V., and Viollier, P.H. (2010). Protein localization and dynamics within a bacterial organelle. Proc Natl Acad Sci U S A *107*, 5599-5604.

S4. Jiang, C., Brown, P.J., Ducret, A., and Brun, Y.V. (2014). Sequential evolution of bacterial morphology by co-option of a developmental regulator. Nature *506*, 489-493.

S5. Radhakrishnan, S.K., Thanbichler, M., and Viollier, P.H. (2008). The dynamic interplay between a cell fate determinant and a lysozyme homolog drives the asymmetric division cycle of *Caulobacter crescentus*. Genes Dev *22*, 212-225.

S6. Randich, A.M., Kysela, D.T., Morlot, C., and Brun, Y.V. (2019). Origin of a Core Bacterial Gene via Co-option and Detoxification of a Phage Lysin. Curr Biol *29*, 1634-1646 e1636.

S7. Saltikov, C.W., and Newman, D.K. (2003). Genetic identification of a respiratory arsenate reductase. Proc Natl Acad Sci U S A *100*, 10983-10988.
